# Recombination impacts damaging and disease mutation accumulation in human populations

**DOI:** 10.1101/006064

**Authors:** Julie G. Hussin, Alan Hodgkinson, Youssef Idaghdour, Jean-Christophe Grenier, Jean-Philippe Goulet, Elias Gbeha, Elodie Hip-Ki, Philip Awadalla

## Abstract

Many decades of theory have demonstrated that in non-recombining systems, slightly deleterious mutations accumulate non-reversibly^1^, potentially driving the extinction of many asexual species. Non-recombining chromosomes in sexual organisms are thought to have degenerated in a similar fashion^2^, however it is not clear the extent to which these processes operate along recombining chromosomes with highly variable rates of crossing over. Using high coverage sequencing data from over 1400 individuals in The 1000 Genomes and CARTaGENE projects, we show that recombination rate modulates the genomic distribution of putatively deleterious variants across the entire human genome. We find that exons in regions of low recombination are significantly enriched for deleterious and disease variants, a signature that varies in strength across worldwide human populations with different demographic histories. As low recombining regions are enriched for highly conserved genes with essential cellular functions, and show an excess of mutations with demonstrated effect on health, this phenomenon likely affects disease susceptibility in humans.

---

The recent human demographic expansion has resulted in an excess of rare variants ^3, 4^, a large proportion of which are putatively functional. Although these variants potentially have phenotypic effects, their distribution across the genome has yet to be fully characterized. Recombination (or linkage) is an important factor in determining the spatial distribution of these rare and potentially disease associated variants along the genome. In the absence of recombination, theory predicts that mildly disadvantageous mutations accumulate on haplotypes, as mutation-free haplotypes cannot be regenerated once they are lost - a process termed “Muller’s ratchet” ^1,5^. Although mostly explored in asexual systems, evidence for this process has recently been described in non-recombining regions of model organisms such as in *Drosophila melanogaster* ^6,7^ or, for mammalian Y-chromosomes, where the suppression of recombination is the primary model explaining why most Y-linked genes are no longer active ^2^, after an irreversible accumulation of genetic defects during millions of years.

Most of the human genome undergoes recombination, at a rate that varies dramatically between different genomic regions; large regions with low recombination called coldspots (CS) are punctuated by short hotspots of recombination where most of the crossovers occur, forming highly recombining regions (HRR) (Figure 1, online supporting note 1). Autosomal coldspots span about a third of the human genome, and 25.3Mb of the exome, while 634.2 Mb genome-wide and 17.8Mb of the exome are within HRRs (Material and Methods, Table S1A). The absolute number of variants within CS is greater than the absolute number of mutations in HRR, although the number of variants per Kb is smaller in CS (Table S1), reflecting reduced neutral genetic diversity in low recombining regions (Figure S1) ^8,9^. Physical linkage of neutral variants to adaptive mutations, “selective sweeps”, are likely to have a role in reducing this genetic variation ^10,11^, but background selection appears to be largely responsible for removal of neutral diversity linked to deleterious variants ^11–13^. Functional variants may also have reduced frequency due to more complex processes, as these alleles can influence each other’s evolutionary dynamics by competing with each other to become fixed within a population ^14^, a process known as Hill-Robertson interference ^1,15^. Conversely, genetic recombination reduces interference by allowing these sites to segregate independently and generate new haplotypes, with the net effect of slowing down the accumulation of rare deleterious variants within exons ^16^.

**Figure 1.**
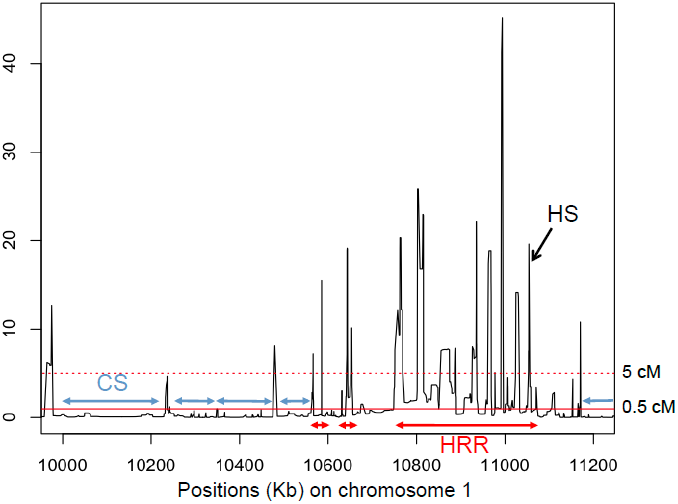
Coldspots (CS). Hotspots (HS) and High Recombination Regions (HRR). CS are defined as regions of more than 50 Kb with recombination rates between adjacent SNPs below 0.5 cM/Mb. HS are defined as a short segment (<15Kb) with recombination rates above 5 cM/Mb. HRR are regions with a high density of hotspots, such that the distance separating neighbouring hotspots (>5 cM/Mb) is smaller than 50 Kb. CS and HRR have to be conserved across FCQ, CEU and YRI genetic maps and to have consistent recombination rates in admixture-based and pedigree genetic maps to be kept in our study (online supporting note 1).

The impact of recombination on diversity via Hill-Robertson interference is a long-standing question among evolutionary biologists, however the theoretical expectations have yet to be empirically demonstrated in human autosomes. Here, we ask whether variable recombination across the human genome has a significant impact on the spatial distribution of deleterious variation. Using single nucleotide variants (SNVs) from RNA and exome-sequencing data we generated for 521 French-Canadians (online supporting note 2) recruited by the CARTaGENE Project ^17,18^ and from high-coverage exome data in 911 individuals from worldwide populations from the 1000 Genomes Project ^19^, we test whether putatively damaging variants accumulate differentially between CS and HRR. Annotation of all SNVs using public resources (Material and Methods) was done using five properties of mutations: the change in amino-acid sequences (non-synonymous or silent variants), the predicted impact on protein function and structure (damaging variants), the level of conservation at the mutated coding site (constrained variants), the allele frequency (rare variants with minor allele frequency lower than 0.01) and the specificity of the variant within populations (private variants). For each category, we quantified the differential mutational burden between CS and HRR using odds ratios (ORs) to assess whether the fractions of polymorphic sites in the given category are significantly different between CS and HRR (Material and Methods).

For all populations, coldspots in the human genome show a higher proportion of rare and non-synonymous variants relative to regions of high recombination (Figure 2, Figure S2-S3). Rare and non-synonymous variants are enriched for mutations of functional importance, with potentially deleterious effects. The site frequency spectrum differs between CS and HRR and minor allele frequencies (MAF) correlate positively with recombination rates (Table S2) ^11^. However, the excess of non-synonymous and damaging variants remains after correcting for MAF (Figure S3). In turn, common synonymous variants at unconstrained (neutral) sites are consistently enriched in HRR compared to CS in all populations. These findings suggest that purifying selection is more efficiently removing harmful variants in high recombining regions, whereas deleterious mutations survive in greater proportion when recombination frequencies are low. We verified that GC/CpG content does not explain the significant enrichment of putatively deleterious mutations in CS in the human exome (Online Supporting Note 2). Furthermore, the results hold even after correcting for average gene expression, substitution rates, types of mutations, exon size and total SNP density per exon (Table S2-S4). Importantly, the results are robust to a wide array of recombination rate thresholds used to define CS and HRR (Table S5). The effect does not appear to be chromosome specific, and no significant difference is observed for genes closer to telomeres (Table S10). We also find that CS have larger introns than HRR (Online Supporting Note 4), consistent with the efficiency of selection being reduced in low recombining regions ^6,20^.

**Figure 2.**
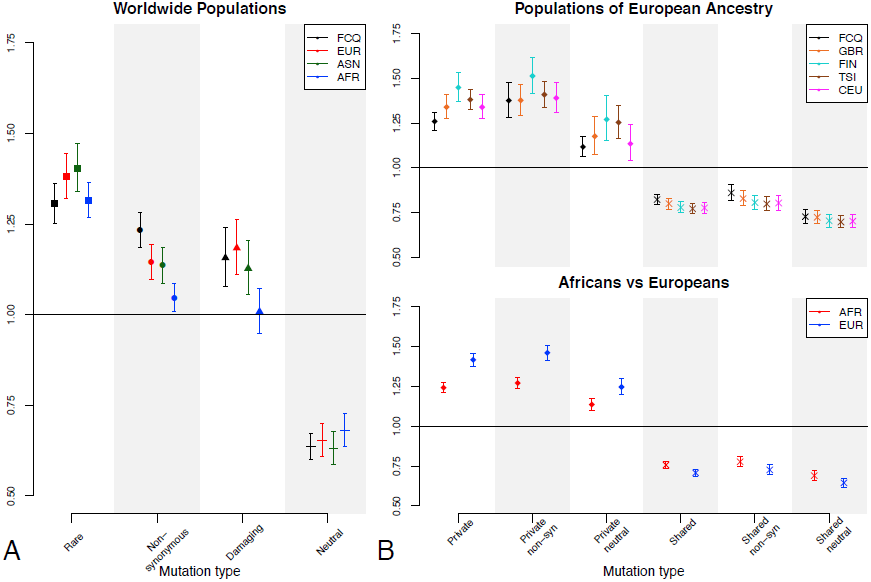
Differential mutational burden in Coldspots (CS) compared to Highly Recombining Regions (HRR) in human populations. Differential burden is computed using odds ratios (OR), representing the relative enrichment of a category of variants compared to all variants in CS versus HRR (OR > 1 means enrichment in CS and OR < 1 means enrichment in HRR). Error bars represent 95% confidence intervals. Variants are categorized as rare (MAF < 0.01 in a population), non-synonymous (missense and nonsense), damaging (predicted by both SIFT and Polyphen2), neutral (common and synonymous), private (specific to a population) and shared (present across populations). (A) Comparison of worldwide populations. ORs are computed based on variants called in exons from 521 French-Canadians of Quebec (FCQ), 379 Europeans (EUR), 286 Asians (ASN) and 246 Africans (AFR). (B) Effects for private and shared variants between populations. Top panel: comparison of closely related populations of West European ancestry. ORs are computed based on private and shared variants called in 96 French-Canadians of Quebec (FCQ), 89 British individuals (GBR), 93 Finns (FIN), 98 Italians from Tuscany (TSI) and 85 European Americans (CEU). Bottom panel: comparison of AFR and EUR populations. Shared variants are present in all populations analysed. Results for different subsets of genomic data are presented in Figure S2 and S3. Results for African sub-populations are shown in Figure S4.

Next, we performed a series of analyses to test whether the genomic spatial distribution of deleterious variants is being modulated through variable Hill-Robertson interference effects in different recombination environments. First, we expect the extent of selective interference to increase with the number of negatively selected sites ^14^, since this would lead to a higher number of interacting alleles. Therefore, we distributed exons into four uniform subgroups based on among-species conservation levels (Material and Methods), which allow for the comparison of exons with similar putative strength of selection in CS and HRR. On average, CS are more conserved than HRR (Figure S5), and binning by conservation levels also ensures that our findings are not influenced by the heterogeneity of conservation across CS and HRR. Conservation levels are determined using the conservation score GERP ^21^ that quantifies position-specific constraint based on the null hypothesis of neutral evolution. The enrichment of rare and putatively deleterious mutations is seen in all conservation classes, showing that even when conservation is normalised across cold and hot regions, exons in coldspots are accumulating more deleterious variants. Moreover, the effect is significantly reduced in classes of exons with lower conservation scores (Figure 3A, Figure S6), as these exons likely have reduced numbers of negatively selected variants interacting with each other. These observations strongly support that selective interference between deleterious variants underlies the observed enrichment of rare and non-synonymous variants in CS compared to HRR, with the results being robust to heterogeneity of selection pressures across the genome.

**Figure 3.**
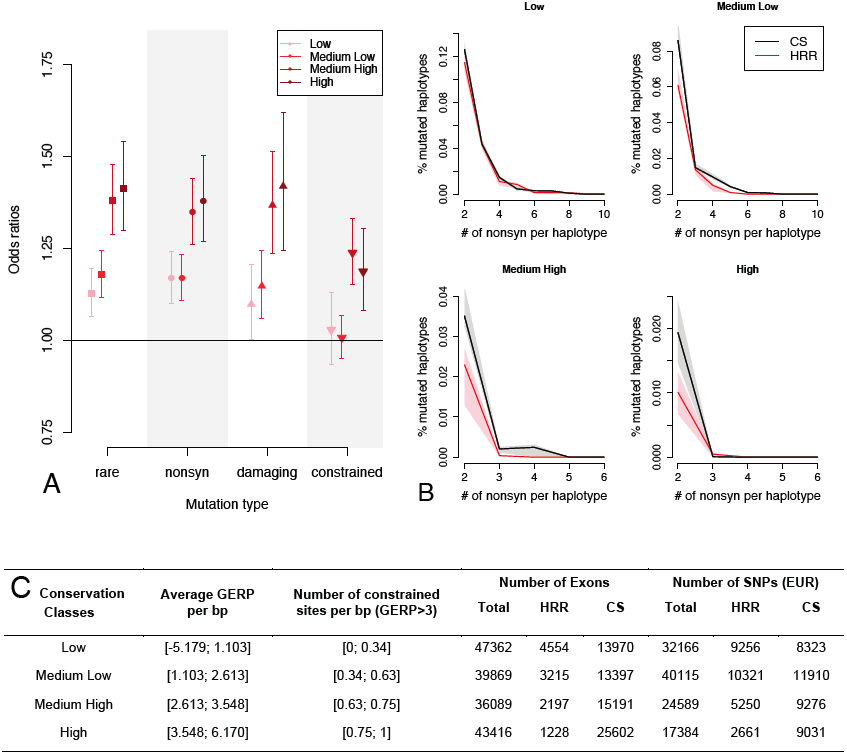
Differential mutational and haplotype burden as a function of conservation. Differential mutational burden between Coldspots (CS) and highly recombining regions (HRR) in Europeans (EUR) measured by Odds ratios for rare (MAF < 0.01), non-synonymous (nonsyn), constrained (GERP < 3 and PhyloP < 1) and damaging variants, for exons binned in four conservation categories (Figure S5). Error bars represent 95% confidence intervals. (B) Haplotype load of non-synonymous variants in CS and HRR. For each conservation category, the proportion of mutated haplotypes with 2, 3, 4, etc. non-synonymous variants in Europeans (EUR) is plotted. The remaining proportion of mutated haplotypes with 1 variant (not plotted) is always larger in HRR than in CS. Confidence intervals are computed by resampling (Material and Methods). The proportion of mutated haplotypes in haplotype class 2 (haplotypes carrying two non-synonymous mutations) is significantly different between CS and HRR, except for the least conserved exons (C) Characteristics of exons in the four conservation categories in terms of average GERP score per base pair (bp) and number of constrained sites per bp (GERP > 3). The number of exons and SNPs considered in plots presented in A and B are reported. Other populations show similar results (Figure S6, S7).

We then evaluated whether haplotypes within human exons have a higher mutational burden in CS compared to HRR using phased data from the 1000 Genomes Project. According to theory and simulations ^1,22,23^, deleterious variants will accumulate faster in low recombination regions in haploid systems, leading to an increased mutational load along individual haplotypes. We computed distributions of non-synonymous variants on haplotypes (Material and Methods), taking the exons as the basic unit, separating them into conservation categories such that compared exons exhibit similar levels of purifying selection. As expected, highly conserved exons tolerate a smaller number of non-synonymous mutations than exons with low conservation. Although within HRR a larger proportion of haplotypes carry at least one mutation compared to CS, because variants are more common and exons are larger (Online Supporting Note 4), haplotypes with two and more non-synonymous mutations are found in significantly higher proportions in CS (Figure 3B, Figure S7). This effect is not seen for the least conserved exons, which are unlikely to be subject to high levels of selective interference. Conversely, we observe a ∼2-fold significant enrichment of the haplotype class with two non-synonymous variants in CS relative to HRR for the most conserved exons, where interference is likely stronger. Although statistical phasing remains technically challenging for rare variants, we see that phasing errors tend to even out rare variants across haplotypes and are unlikely to drive the effect seen here, as the observed bias is similar between CS and HRR (Figure S12, Online Supporting Note 3). This differential haplotype burden is in line with a Muller’s ratchet process where the least-loaded haplotype class, eroded by drift or mutation, is not replenished as fast when there is a lack of recombination, causing haplotypes with more than one deleterious variant to be slightly more frequent.

We also performed extensive computer simulations (Online Supporting Note 3, Table S6) using forward-in-time simulators ^24,25^ to determine whether the above observations are consistent with interference between negatively selected mutations in diploid genomes with variable recombination rates. Simulated coldspots show a higher proportion of rare and negatively selected mutations on haplotypes compared to HRR under a number of different demographic models (Table S7). The differences between recombination environments increase with the proportion of selected sites simulated and we find that a small proportion of strongly selected mutations are less impactful than many mutations with small effects. The effects of background selection are also predicted to be stronger under similar conditions ^26^. However, models of background selection without interference between deleterious sites do not show an enrichment of rare neutral mutations in CS compared to HRR (Figure 4). These simulation results strongly support that linkage between mildly deleterious sites is required to generate the patterns of diversity we have observed. Thus, the effects on neutral diversity attributed to background selection from independent deleterious mutations are likely reinforced by selective interference between these weakly selected mutations. Fixed mutations under negative selection show no significant enrichment in simulated coldspots, indicating that selective interference within populations does not necessarily result in higher rates of fixation of deleterious mutations in diploid organisms ^27^, thereby causing inter-species divergence measures to be insensitive to these effects ^28^. Altogether, these analyses present clear evidence that selective interference is a determining factor shaping patterns of diversity along human autosomes, although other processes, such as heterogeneous mutation rates and variable selection pressures within exons, may contribute, at least in part, to the observed pattern.

**Figure 4.**
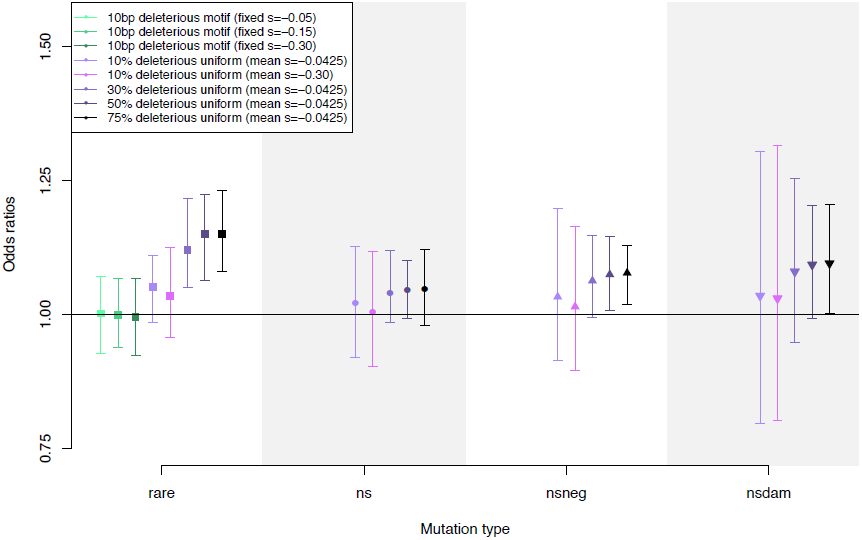
Simulations under different background selection models, with and without interference. Differential mutational burden is computed using odds ratios (OR), representing the relative enrichment of a category of variants compared to all variants in simulated CS vs. HRR. Error bars represent 95% confidence intervals. Non-synonymous (ns) mutations are site attributed a negative selection coefficient and were divided in 3 categories: non-synonymous with a selection coefficient larger than -1/N_e_ (*ns*, effectively neutral), smaller than -1/N_e_ (*nsneg*, negatively selected) and smaller than -0.01 (*nsdam*, damaging). The models with a single locus under negative selection (10 bp deleterious motif, green) are background selection models without linkage between negatively selected sites. The other models have the same parameters as the EW model with µ=r (parameters in Table S6) and include linkage between different proportion *p* of sites under negative selection. Online Supporting Note 3.3 includes further description of background selection models.

If selection is driving this process, we may expect to see variable effects in populations with different histories and effective population sizes. To compare effects across populations, we considered variants in highly covered exons across all datasets (Table S1). We found a relative enrichment for non-synonymous variants in CS compared to HRR in all populations (Figure 2A, Figure S2), but the effect is reduced in African populations compared to others. Furthermore, damaging variants are enriched in CS for French-Canadian, European and Asian populations but not for Africans, making this effect potentially modulated by demographic history. Enrichment of variants private to Africans and Europeans in CS is significantly stronger in Europeans (Figure 2B) even after correcting for MAF (Figure S3C-D), whereas shared variants are more equally under-represented in CS, suggesting that population history impacts the distribution of novel deleterious mutations. Population-specific variants in the five relatively recently diverged populations of European ancestry are also enriched in CS compared to HRR. Interestingly, this effect is also observed for variants that are private to the recently founded French-Canadian population, even after adjusting for GC-content and total SNP density (Table S2). These mutations were either very rare in the source population, or they originated *de novo* since the founding of Quebec four hundreds years ago ^29^. The potential reduction in the efficiency of purifying selection in coldspots is thus detectable over brief evolutionary time scales and suggests that selection may have been affecting the distribution of variants differentially along the genome in the recent past.

Finally, for each individual we computed the relative proportion of rare and non-synonymous variants in CS and HRR and the resulting odds ratios (ORs)(Figure 5, Figure S8). The distribution of per-individual ORs reveals extensive differences between individuals and populations. Although at the population level, rare and non-synonymous variants are enriched in CS compared to HRR (Figure 2A), these effects are not observed at the individual level in Africans. For rare variants, Europeans and Asians all have significant ORs (Figure S8), but the mean individual OR is larger in Europeans. Strikingly, the FCQ population shows an increased variance relative to other populations and exhibits more extreme OR values. For non-synonymous variants, few individuals among AFR, ASN and EUR exhibit significant ORs, whereas again, the FCQ distribution is remarkably different and shows a shift in mean towards larger and significant individual effects (Figure 4B). For private variants, French-Canadians are the most extreme of European populations, followed by Finns and Italians from Tuscany (Figure S9). To confirm these results, we sequenced the exomes of 96 of the FCQ individuals, which fully support the differences seen in this population (Figure S10). We also verified that these differences are not due to discrepancies in SNP annotations and ascertainment between cohorts (Online Supporting Note 4, Figure S11). Average recombination rates within CS and HRR differ between Africans and Europeans (Online Supporting Note 1), however, our simulations show that this heterogeneity in recombination rate is unlikely to be sufficient to explain the observed patterns (Figure S12, Online Supporting Note 3). Differences in population sizes and demographic histories could impact selective interference and variation in CS more than in HRR as selection is less efficient in smaller populations ^29^. The population-specific increase in the differential burden observed here may be due to complex recent demographic events and depart from Wright-Fisher’s expectations (Online Supplementary Note 3), such as serial founder effects or range expansions ^30^. Therefore, complex demographic processes in humans may have impacted genetic variation and the effective strength of selection differentially along the genome through the history of the human peopling. Although studies suggest that recent population history has little impact on the burden of deleterious mutations ^31^, our study questions the generality of these findings and documents another way in which population history can influence patterns of deleterious genetic variation across populations.

**Figure 5.**
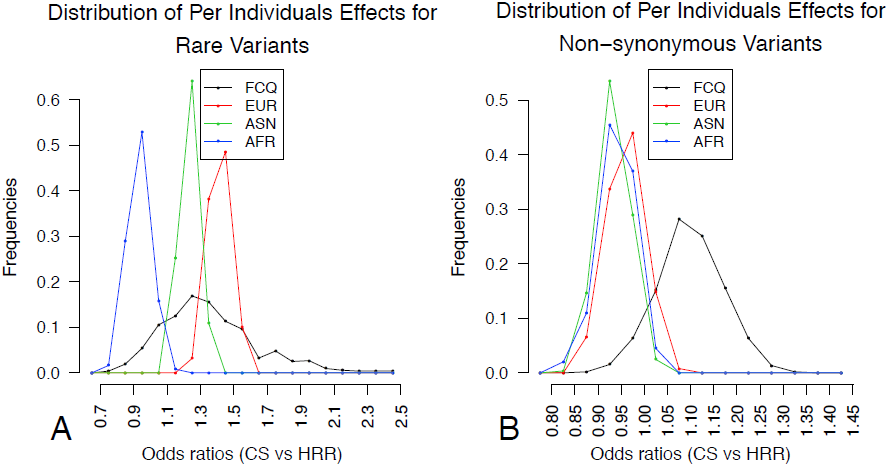
Per individual differential mutational burden across populations. Distribution of odds ratios (OR) per individual in French-Canadians, Europeans, Asians and Africans, comparing proportions of (A) rare and (B) non-synonymous variants between coldspots (CS) and high recombining regions (HRR). The frequencies of individual OR in each population are shown. Distributions are significantly different between populations for all comparisons (Kruskal-Wallis test, p < 5e-9) except for AFR versus ASN non-synonymous OR distributions (Kruskal-Wallis test, p = 0.051). The relative proportions of rare or non-synonymous mutations in CS and HRR are shown in Figure S8. Distributions for private variants in European sub-populations are shown in Figure S9. French-Canadian data used is the RNAseq dataset (Online Supporting Note 2), replication with exome data of 96 French-Canadians is presented Figure S10. These results are robust to annotation pipeline and exclusion of fixed polymorphisms (Figure S12).

The irreversible accumulation of mildly deleterious mutations along haplotypes in low recombining regions can have damaging phenotypic effects in individuals carrying them, since genomic regions with high linkage disequilibrium show an excess of genes primordial for response to DNA repair and cell cycle progression ^32.^ Furthermore, CS are enriched for genes involved in protein metabolic processes, mRNA processing, organelle organisation, microtubule-based processes and genes highly mutated in cancer (Table S8-S9, Online Supplementary Note 4). Selective interference between deleterious mutations may thus impact the genetic aetiology of human diseases. By examining sequence variants that have a demonstrated effect on health reported in the ClinVar database ^33^, we found that for variants with MAF lower than 0.01 in humans, disease-related mutations reported in ClinVar are enriched in CS relative to HRR (Table S9). The effect is driven mainly by the coldest recombination regions in the human genome (mean rate lower than 0.1 cM/Mb) and decreases with the increasing recombination rate (Figure S6), which is expected under selective interference and cannot be explained by ascertainment bias within ClinVar (Online Supplementary Note 4). Ultra-sensitive regions of the genome, where a 400-fold enrichment of disease-causing mutations has previously been reported ^34^, are greatly enriched in CS (Figure 6, Table S9). Altogether, these results indicate that rare variants identified as medically relevant are more likely to be found in human coldspots, as new disease-causing mutations occurring in regions with higher recombination rates will be more efficiently eliminated by natural selection.

**Figure 6.**
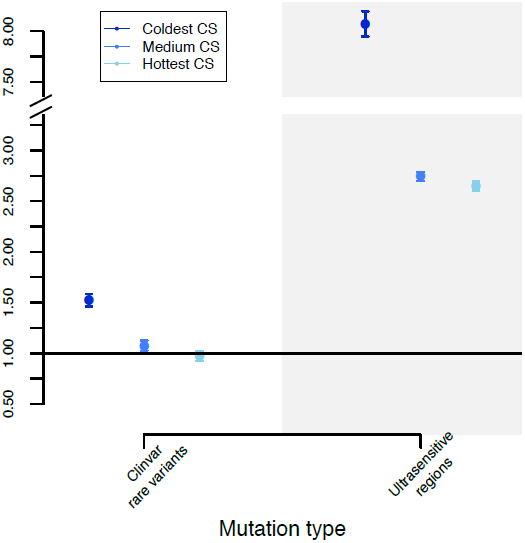
Differential mutational burden for rare disease mutations and sensitive regions. Relative enrichment of rare disease mutations listed in the ClinVar database and in Khurana et al. 2013 ^34^ for ultra-sensitive motifs in CS relative to HRR, in coding regions only. Error bars represent 95% confidence intervals. CS regions are binned into 3 categories such that each group roughly contains the same number of regions: coldest (recombination rate *r* <0.1 cM/Mb), medium (0.1 ≤ *r* <0.15 cM/Mb) and hottest (0.15 ≤ *r* <0.5 cM/Mb). Rare disease mutations have MAF < 0.01 or are not segregating in the 1000 Genomes populations. In each category of CS, enrichment of rare disease mutations is evaluated by comparison to the number of rare variants in the 1000 Genomes populations. Enrichment for ultra-sensitive motifs is evaluated by correcting for sequence length in each category of CS.

Until now, selective interference had been empirically studied through inter-species divergence among primates, and no effect of recombination on such measures had been detected ^28^. In contrast, our population genetic analyses reveal that the efficiency of selection depends on local recombination rates modulating selective interference, explaining the spectrum of rare functional variation. Furthermore, recent population demography likely impacts the differential mutational burden, with selective interference potentially modulating the accumulation of new deleterious mutations more strongly in small bottlenecked populations. The spatial distribution of deleterious mutations along the genome would therefore vary according to recent population history. Finally, features such as recombination rates, driving genetic variation heterogeneity across the genomic landscape, have a potential impact on disease mutation accumulation and disease mapping strategies. Our deeper understanding of how the processes of recombination, selection and mutation work together to shape the landscape of deleterious diversity in the genome will thus improve our ability to map disease-causing mutations in humans.

## Material and Methods

### Genetic maps and recombination

We used three LD-based genetic maps ^35^, one pedigree map ^36^ and one admixture-based map ^37^ to identify ‘cold’ or ‘hot’ regions in the human genome that were shared across all human populations (Figure 1). The three LD-based maps were built from genotyping data from French-Canadians from Quebec (FCQ), CEU and YRI from HapMap3 (see Online Supporting Note 1). We used these maps to locate coldspots and hotspots of recombination. We define coldspots (CS) as regions of more than 50Kb with recombination rates between adjacent SNPs below 0.5 cM/Mb in FCQ, CEU and YRI populations. A hotspot is defined as a short segment (<15Kb) with recombination rates falling in the 90th percentile (> 5 cM/Mb). We define high recombination regions (HRR) as regions with a high density of hotspots, such that the distance separating neighbouring hotspots is smaller than 50 Kb. For each CS and HRR identified, we computed the mean recombination rate (cM/Mb) using deCODE pedigree-based map and the admixture-based African American map, and excluded all regions with inconsistent recombination rates in one of these maps. We obtained a list of 7,381 autosomal coldspots, spanning about a third of the human genome, for a total of 1.049 Gb (Table S1A). We identified 12,500 HRRs genome wide shared between FCQ, CEU and YRI populations, covering a total of 634.2 Mb. We verified that our results are robust to the choice in recombination parameters used to define regions (Table S5) and to differences between LD-based and pedigree maps (Online Supporting Note 4).

### Genomic data

Genomic data from the 1000 Genome Project and the CARTaGENE project were used in this study. SNP calls for 911 individuals from 11 non-admixed populations from the 1000 Genomes Project Phase 1 high coverage exon-targeted data (50-100✕) were downloaded from the 1000 genomes ftp site. Only SNPs called within targeted exons were extracted from vcf files. Details on 1000 Genomes populations, sequencing protocol, SNP calling, and validation can be found in the 1000 Genomes phase 1 publication ^19^. Populations were then grouped by continental ancestry, with a total of 142,296 SNPs in 379 Europeans (EUR), 128,697 SNPs in 286 Asians (ASN) and 186,549 SNPs in 246 Africans (AFR) within 124,015 different exons (AFR populations: ASW, YRI, LWK; EUR populations: FIN, TSI, IBS, GBR, CEU; ASN populations: CHB, CHS, JPT). Admixed American populations (MXL, CLM and PUR) were excluded from analyses. For the French-Canadian population, SNPs were called from RNA sequencing data (RNAseq), with a total of 178,394 SNPs in 521 individuals within expressed exons. RNAseq 100 bp pair-ends indexed libraries were constructed using the TruSeq RNASeq library kit (Illumina). Sequencing was done on HiSeq machines (Illumina), multiplexing three samples per lane. More details on the samples, the sequencing protocol and downstream analyses are described in supporting online note 2 and in ^18^. Analyses comparing populations were done on a subset of exons that are highly covered (> 20✕, HC exons) across all datasets. A total of 89,390 exons passed this filter, containing a total of 73,627 SNPs in FCQ, 69,672 in EUR, 63,726 in ASN and 89,789 in AFR (Table S1B). Additionally, all FCQ individuals were genotyped on the Illumina Omni2.5M array, and a subset of them (96 individuals) was exome sequenced on HiSeq machines (Illumina) with a total of 60,251 SNPs called (online supporting note 2).

### Variant and exon annotations

We annotated SNPs either as synonymous or non-synonymous and those reported as intronic, in untranslated regions (UTR) or in non-coding RNA were labeled as ‘other’. Details on annotation pipelines are given in Online Supporting Note 4.1. Functional annotations for non-synonymous mutations were obtained with PolyPhen2 and SIFT ^38,39^ and a variant was annotated as damaging when both methods predict the mutation to be damaging. Because there is a bias in these methods towards seeing a reference allele as benign, we excluded variants where the major allele was not the reference allele. Nonsense variants were annotated as damaging. We retrieved GERP conservation scores for all positions within the human exome (mendel.stanford.edu/SidowLab/downloads/gerp/index.html) and PhyloP conservation scores from UCSC Genome website (hgdownload.cse.ucsc.edu/goldenpath/hg19/phyloP46way). We annotated a mutation as constrained if GERP > 3 and PhyloP > 1. Minor allele frequencies (MAF) were computed for each population independently and rare variants herein have a MAF < within a population. SNPs with MAF > 0.01 are annotated as ‘neutral’ if they are synonymous and have both GERP < 3 and PhyloP < 1. For European populations, including FCQ, we annotated SNPs as private when they were unique to a population (not in other European populations nor in ASN and AFR) whereas shared variants are SNPs in common between all European populations. We also annotated exons by tabulating exon size, GC-content, average recombination rates in FCQ, CEU and YRI population genetic maps, average gene expression (computed from the FCQ RNAseq data, online supplementary note 2), average GERP score, number of sites with GERP > 3, rate of synonymous substitutions per site (*dS*) between homologous gene sequences from human and chimp. *dS* values were retrieved from Ensembl as of November 2013, for 15,075 orthologous genes out of the 21,986 RefSeq genes and have been calculated by *codeml* in the PAML package ^40^. Exons were ranked based on GC-content, GERP annotations, *dS* values and expression level, and stratified in four categories of equal sizes: low (25% lowest values), medium low (ranked between 25 and 50%), medium high (ranked between 50 and 75%) and high (25% highest values). For each exon and each population, we computed the number of SNPs, average MAF per population and the density (SNP/Kb) of non-synonymous, constrained and damaging variants.

## Differential mutational burden

Odds Ratios (OR) are used to compare proportion of mutations between CS and HRR, for different types of mutations, and we call this measure of enrichment the differential mutational burden. The variant types analysed are *rare, non-synonymous, damaging, constrained, neutral, private, non-synonymous private* and *shared*. The OR for a variant type T is computed as:

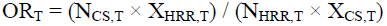

with N_CS,T_ and N_HRR,T_ the number of variants annotated of type T and X_CS,T_ and X_HRR,T_ the number of other annotated variants in CS and HRR, respectively. Variants with missing annotations for a given type were excluded of the OR calculation for that type. Confidence intervals were calculated following the procedure described in ^41^. Using alleles carried by each individual at all annotated variants, the differential mutational burden is also evaluated per individual. We consider zygosity per site, such that for a heterozygote at a non-synonymous site for example, both the non-synonymous allele and the reference allele are counted, and for a non-reference homozygote, the non-synonymous site is counted twice. We note that ignoring zygosity (counting each site once as either mutated or reference) does not change the distributions of individual ORs. Sites for which the reference allele is the derived allele (as inferred from chimp and rhesus macaque) were ignored in the per-individual OR calculation. We evaluated correlations between recombination rates and variant density in exons using linear regression models, adjusting for expression level (online supporting note 2), GC-content, exon size and SNP density.

### Haplotype load

Haplotypes from the 1000 Genomes populations were phased using ShapeIT2 using SNP array and sequence data (ftp://ftp.1000genomes.ebi.ac.uk/vol1/ftp/release/20110521/) as of November 2013 ^42^. For each phased haplotype in an exon, we counted the number of non-synonymous mutations, and we computed haplotypes allelic classes ^43^, ie. the number of haplotypes with 0, 1, 2, …, n_max_ non-synonymous mutations. We identified 9 genes with exons that have an extremely high haplotype load: HRNR, FLG, MKI67, OR51B6, AHNAK2, PLIN4, CMYA5, MUC17, RP1L1. These genes were excluded from the analyses. After excluding these outlier exons, the maximum number n_max_ of non-synonymous mutations on a haplotype never exceeds 10. CS and HRR haplotype distribution were compared for exons in each conservation category. Conservation categories are based on a conservative ranking of exons according to both their average GERP scores and their number of sites with GERP > 3 (Figure S5). Confidence intervals for the proportions of haplotypes in each haplotype class are computed by resampling the same number of exons in CS and HRR in each conservation category.

### Simulations

We perform forward simulations using SLiM and sfs_code ^24,25^. For 250 diploid individuals (500 haplotypes), we simulated exon-like sequences with non-synonymous mutations, with the mutation rate µ = 2×10^-7^ per base and N_e_ = 1000, and tested models with the overall recombination rate being r = µ or r = µ/2. We simulated human-like recombination environments, according to the average recombination rate and size of CS and HRR in the human exome: CS are 95 Kb long in average and contain 4.1% of recombination, HRR are 28 Kb long in average with 58.6% of recombination, according to values in the deCODE map. We also considered CS and HRR with different recombination rates, to match the African recombination rates (Online Supporting Note 3). Each simulated exome consists of 75 fragments of 200 Kb, with CS- and HRR-like sequences separated by 77Kb. Different demographic and selection models are studied (Online Supporting Note 3) with parameters inferred in previous studies ^44,45^. The fitness effects for non-synonymous mutations are gamma distributed with mean *s,* with dominance coefficient *h* = 0.5. We ignored positive selection in our models. We also performed simulations under null models, with constant r or without negative selection (*s* = 0), to evaluate expectations without variation in recombination rates or under selective neutrality. For each simulation, mutations that have a derived allele frequency below 0.01 were annotated as *rare*. Non-synonymous mutations are site attributed a negative selection coefficient and were divided in 3 categories: non-synonymous with a selection coefficient larger than -1/N_e_ (*ns*, effectively neutral), smaller than - 1/N_e_ (*nsneg*, negatively selected) and smaller than -0.01 (*nsdam*, damaging). The threshold for damaging variants is set to match the number of non-synonymous damaging variants in the empirical data. We used odds ratios to estimate the differential mutational burden between CS and HRR for rare and non-synonymous mutations in simulations.

### Disease mutations and genes

Clinical variants with potential and validated pathogenicity were retrieved from the curated ClinVar database ^33^ (https://www.ncbi.nlm.nih.gov/clinvar). Variants that are not segregating 1000 Genomes populations (which represent the vast majority of ClinVar variants), or that are found at a MAF lower than 0.01 in these populations, were classified as *rare*. ORs were computed by comparing number of rare variants to non-disease variants with MAF in 1000 Genomes populations lower than 0.01 between CS and HRR. Sensitive and ultra-sensitive regions coordinates were retrieved from ^34^, that have been examined for the presence of inherited disease-causing mutations from the Human Gene Mutation Database (HGMD). We used PANTHER ^46^ and WebGestalt ^47^ resources to perform gene set enrichment analyses (Online Supporting Note 4).

#### Contributions and Acknowledgments

JH, AH, and PA designed the study. PA provided samples. EG and EHK processed samples for sequencing and genotyping. JCG and VB aligned sequencing data. JH, YI, AH, JCG, and JPG performed quality control on genotyping and sequencing data. JH performed the analyses. JH, AH, and PA wrote the manuscript. We thank Claude Bhérer, Greg Gibson, Edouard Hatton, Gil McVean, John Novembre, Chris Spencer, Eric Stone and anonymous reviewers for insightful comments on the study, and the CARTaGENE participants and team for data collection. We confirm that informed consent was obtained from all subjects. We acknowledge financial support from Fonds de la recherche en santé du Québec (FRSQ), Genome Quebec, Fonds québécois de la recherche sur la nature et les technologies (FQRNT) and the Canadian Partnership Against Cancer. JH is a Human Frontiers Postdoctoral Fellow, AH is an FRSQ Research Fellow and YI is a Banting Postdoctoral Fellow. The authors declare no conflict of interests. Requests for CARTaGENE data published here should be submitted to the CARTaGENE program at, citing this study. Other datasets are available from www.well.ox.ac.uk/∼julieh/mshussin2014

## Online Supporting Information

### Online supporting note 1– Recombination analyses

#### 1. Overview

Accurate population recombination maps are necessary to identify regions that are in different recombination environments. We used five population genetic maps (*1-3*), to identify ‘cold’ or ‘hot’ regions. We first identified these regions using LD-based population maps, and in a second step, we excluded regions for which recombination rates in pedigree and admixture maps were inconsistent with the definitions of the cold and hot regions. The reason to first consider LD-based methods is that, we wanted to identify regions that have been low recombining (or high recombining) for a considerable amount of time during human evolution, as selective interference is a phenomenon that would occur over many generations. Pedigree map give infomation on the recombination rates in the current generations, which does not guarantee that a given region was not recombining in the evolutionary past of humans. LD-based methods, although potentially biaised by SNP density, by using polymorphism data from population samples, look at ancestral recombination rates and provide us with the opportunity to identify regions with no evidence of recombination over hundred of thousands of years.

#### 2. Population genetic map of French-Canadians

We build the genetic map of the French-Canadians of Quebec (FCQ) population using LDhat (*4*). The 521 FCQ individuals were genotyped on the Illumina Omni2.5M array. A total of 1,554,440 autosomal SNPs were obtained after filtering (Quality control HWE p<0.001, Missingness < 0.05, MAF > 0). We ran the *interval* program from the LDhat package on FCQ genotyping data. Because the likelihood tables for the *interval* program are pre-computed for a maximum number of 192 haplotypes, we randomly selected 96 unrelated individuals from the 521 FCQ individuals. The largest chromosomes (1 to 12) were broken into two segments (p and q arms) and all genomic segments were phased with ShapeIT2 (*5*). We ran the *interval* program on each genomic segment for 30,300,000 iterations with a burn-in of 300,000 iterations and sampled the population recombination rates ρ every 10,000 iterations. The estimate of the recombination rate ρ between each pair of adjacent SNPs, in units of 4N_e_r per Kb, was computed by taking the average rate across iterations of the rjMCMC procedure implemented in *interval*. We sampled the population recombination rates ρ every 10,000 iterations. To convert the population recombination rate estimates in 4N_e_r per Kb into centiMorgan per Megabase (cM/Mb), we inferred the effective population size N_e_ for the FC population using the estimates of r computed for the 2010 deCODE map in cM units (*2*). Specifically, we identified chromosomal segments where both FCQ data and deCODE SNP positions allowed estimates of rates and we summed rates across these genomic regions to obtain the total estimated distance (4N_e_R) and the total genetic distance (R in cM units) from the deCODE map.

#### 3. Population genetic maps from HapMap3 data

Population genetic maps from the HapMap2 data have been built by the HapMap consortium in 2007 (*3*), using the 2002 deCODE pedigree map (*6*) and hg18 positions. It was subsequently ‘lifted over’ to hg19 using the UCSC liftOver tool and regions where the order of markers had changed were removed from the final maps. We re-computed the HapMap maps for CEU and YRI with the methodology used to compute the FCQ map described above to allow direct comparison. Specifically, we performed a lift over on the HapMap3 SNPs positions prior to estimating recombination rates with *interval* using 96 unrelated individuals from the CEU and YRI populations. We then converted the recombination rates in cM/Mb using the 2010 deCODE pedigree map (*2*), and obtained new HapMap genetic maps for these populations. These maps are available here: www.well.ox.ac.uk/∼julieh/mshussin2014

#### 4. Coldspots and High Recombining Regions

We used these genetic maps to locate coldspots and hotspots of recombination. We define coldspots (CS) as regions of more than 50Kb with recombination rates between adjacent SNPs below 0.5 cM/Mb in FCQ, CEU and YRI populations, such that they are shared between all human populations studied. We excluded centromeric regions and required that at least 5 SNPs support the coldspot, to avoid regions with dramatically reduced diversity, where power to estimate recombination rates is decreased. For each region identified, we computed the mean recombination rate (cM/Mb) using the deCODE pedigree map (*2*) and the admixture-based African American map (*1*), and we excluded all regions that have a recombination rate larger than 0.5 cM/Mb in one of these maps. We obtained a list of 7,381 autosomal CS, spanning about a third of the human genome, for a total of 1.049 Gb (Table S1). A hotspot is defined as a short segment (<15Kb) with recombination rates falling in the 90th percentile (> 5 cM/Mb). We define high recombination regions (HRR) as regions with a high density of hotspots, such that the distance separating neighbouring hotspots is smaller than 50 Kb. We identified 12,500 HRRs genome wide shared between FC, CEU and YRI populations, covering a total of 634.2 Mb (Table S1). The definition of coldspot, hotspot and HRR are illustrated in Figure 1. A complete list of these regions can be found at www.well.ox.ac.uk/∼julieh/mshussin2014. The recombination rate thresholds used to define coldspots and hotspot were chosen to maximize the overall number of SNPs included in the analyses while minimizing the difference between the number of SNPs in coldpots and in HRRs. The effects are robust to different recombination thresholds (Table S5).

Although these regions are inferred in both YRI and CEU LD maps, they may have different recombination rates. We compared the mean rates per CS and HRR between these two LD-based maps. The mean recombination rate in CS is 0.129 cM/Mb for CEU and 0.181 cM/Mb for YRI, with this difference being highly significant (p < 10^-5^, permutation test). In HRR, the mean recombination rate in CS is 5.70 cM/Mb for CEU and 4.62 cM/Mb for YRI (p < 10^-5^, permutation test). The distributions of rates within these regions are highly different between YRI and CEU (Kruskal-Wallis chi-squared = 943.0448, df = 1, p-value < 2.2e-16). These differences could be due to differences in LD-based maps caused by varying demography and population specific selection; however, it is more likely that it reflects differences in local recombination rates due to the presence of different alleles of PRDM9, the protein responsible for recombination clustering in hotspots along the genome (*1, 7*). The impact of these differences in mean rates in CS and HRR between African and non-African populations is explored in Online Supporting Note 3.5 and Figure S11 and is unlikely to cause the large differences we observe between populations in Figure 5.

#### 5. Comparison of CS and HRR with the deCODE map

Only 481 coldspots were excluded because recombination rates were larger than 0.5 cM/Mb in the deCODE map. The deCODE recombination rate for each CS is reported in the supplementary data available online at www.well.ox.ac.uk/∼julieh/mshussin2014

To ensure our result are robust to the choice of map, we computed CS and HRR using the deCODE pedigree map alone and compared this set of regions to the set of regions used in our study. There are 8165 CS inferred from the deCODE map, and 1824 CS supported by more than 5 SNPs that do not overlap a CS in our final list of coldspots:

- 94 had been removed because of a lack of SNPs in the region in HapMap or FCQ data (less than 5 SNPs);
- 1324 have a recombination rate > 0.5cM/Mb in all LD-based maps;
- For the remaining 406 CS :

○ 121 have recombination rate > 0.5cM/Mb in FCQ LD map;
○ 125 have recombination rate > 0.5cM/Mb in CEU LD map;
○ 142 have recombination rate > 0.5cM/Mb in ASN LD map;
○ 332 have recombination rate > 0.5cM/Mb in YRI LD map.

For HRR, a smaller number of regions are found with the deCODE map (9071). Discordances in HRR lists are mainly due to differences in the computed intensity of recombination hotspots by the two methods, that are not directly comparable.

We recomputed the enrichment statistics for rare, non-synonymous, damaging and neutral variants for the FCQ dataset with the deCODE regions alone. Rare (1.11[1.06;1.16]), non-synonymous (1.10 [1.06;1.15]) and damaging (1.08 [1.01,1.16]) variants remain significantly enriched in CS and neutral variants are significantly underrepresented (0.85 [0.81;0.91]). However, the effects are somewhat weaker, which is explained by the fact that we include CS with evidence of recombination in LD-maps. If we only take the overlap (ie. variants in CS in both deCODE and LD-maps), the effects become comparable to the ones observed in the final list of CS obtained: rare (1.27[1.22;1.33]), non-synonymous (1.17 [1.12;1.22]), damaging (1.05 [1.01,1.11]) neutral (0.68 [0.65;0.73]). Our results are therefore robust to the choice of map.

### Online Supporting Note 2 – French-Canadian Genomic Data

#### 1. Overview

The main analyses for the French-Canadian (FCQ) population rely on SNPs called from RNA sequencing (RNAseq) data. SNPs from other populations come from the 1000 Genomes phase I high coverage exome dataset and were grouped according to their ancestry (African, European, Asian ancestry). Admixed populations from the Americas were excluded. We performed extensive analyses to ensure that calling SNPs from transcriptomes do not create biases influencing our population genetic analyses and cannot explain the difference seen in the FCQ population. We also performed exome sequencing for a subset of FCQ individuals for which we had RNAseq data and further validated SNP calls and the overall results found with the RNAseq SNPs.

#### 2. French Canadians

The CARTaGENE project (CaG) collected biologicals and data from 20,000 participants recruited throughout the province of Quebec (*8*), and high-density genotyping and RNA sequencing data was generated for 521 French-Canadians participants (Material and Methods). Sampling includes individuals from three distinct metropolitan regions of Quebec: the Montreal area (MTL), Quebec City (QCC) and the Saguenay Lac-St-Jean region (SAG) (Figure S13). Regional origins of the individuals were validated with a principal component analysis (PCA) of genetic diversity using genotypic data and including individuals from the Reference Panel of Quebec (RPQ) (*9*). Population structure is complex and made of regionally differentiated populations (Figure S13), resulting from the very recent regional founder effect that occurred in Saguenay. This territory was colonized during the 19th century by a reduced number of settlers, who contributed massively to the genetic pool of individuals living in this region today (*10*).

#### 3. Processing of the raw RNAseq Data and SNP calling

Approximately 3 mL of blood was collected for RNA work in Tempus Blood RNA Tubes (Life Technologies). Total RNA was extracted using a Tempus Spin RNA Isolation kit followed by globin mRNA depletion by using a GLOBINclear-Human kit (Life Technologies). RNAseq 100 bp pair-ends indexed libraries were constructed using the TruSeq RNASeq library kit (Illumina). Sequencing was done on HiSeq machines (Illumina), multiplexing three samples per lane. After initial filtering based on sequencing read quality, paired-end reads were aligned using TopHat (V1.4.0) (*11*) to the hg19 European Major Allele Reference Genome (*12*). PCR removal was performed using Picard (picard_tools/1.56, http://picard.sourceforge.net). Raw gene-level counts data were generated using htseq 0.5.3p3 (*13*). These counts were then normalized using EDASeq v1.4.0 and a procedure that adjust for GC-content as well as for distributional differences between and within sequencing lanes (*14, 15*). Average normalized gene expression levels per gene were determined by averaging expression levels of each gene across all individuals (Idaghdour et al. 2014. *In preparation*). Every exon of a gene was attributed the gene-level value.

SNP were called from RNAseq data using a procedure similar to SNP calling in exome sequencing data. However, prior to SNP calling, bowtie2 (0.12.7)(*16*) was used to removed abundant sequences (polyA, polyT, tRNA). Only reads that were properly paired and uniquely mapped were kept. Mapping quality score were recalibrated using GATK (*17*) and SNP calling was performed with samtools (0.1.18) (*18*). Filtering of SNPs was done using vcftools v0.1.7 (*19*). We kept SNPs with variant quality of 30 and genotype quality of 20 (Phred scores). Minor allele frequencies (MAF), the proportion of individuals with non-missing genotypes and Hardy-Weinberg equilibrium (HWE) p-values were computed using plink v1.07. SNPs showing departures from HWE at *p* < 0.001 were excluded. We obtained a total of 178,394 polymorphic SNPs (MAF > 0) in the 521 French-Canadians individuals (Table S1B).

#### 4. Selection of Highly Covered Exons

To insure that sequencing SNPs are called throughout the length of exons and to reduce the possible biases due to read depth, we selected highly covered exons (hereafter termed HC exons) with all positions of their sequence covered at a minimum of 20✕ in more than 50% of the sequenced individuals (i.e. at least 261 FCQ individuals). We used BAMStats-1.25 to obtain the minimum coverage per exon per individual for 208,226 autosomal exons. A total of 89,390 exons passed this stringent filter containing a total of 73,627 SNPs. For subsequent analyses, we also excluded 9 genes for which the mutational profiles were abnormal (Material and Methods).

#### 5. FCQ Exome Sequencing Data and Genotyping

All individuals were also genotyped on the Illumina Omni2.5M array. A total of 1,554,440 autosomal SNPs were obtained after filtering (Quality control HWE p < 0.001, MAF > 0). We took all positions in common between the Omni2.5 chip and the RNAseq SNPs called. We filtered for missingness (<50% for RNAseq, <95% Omni) and filtered out positions for which the alleles did not match between the chip and RNAseq after flipping, ending up with 26,615 positions to compare. We compared genotypes for which there was a call in both datasets, and the concordance rates were above 98.8% in all individuals, with mean across individuals of 99.3%.

Exome sequencing was also performed for 96 FCQ individuals. DNA from each sample was extracted from peripheral blood cells and paired-end exome sequencing was performed on HiSeq machines (Illumina), multiplexing six samples per lane. We first performed trimming of sequencing read data using Trim Galore prior alignment to trim adaptors (with parameter –q 0, www.bioinformatics.babraham.ac.uk/projects/trim_galore). Alignment was performed using BWA version 0.5.9-r16. After recalibration with GATK (*17*), reads were trimmed for quality using bamUtil version 1.0.2 (genome.sph.umich.edu/wiki/BamUtil). SNP calling was performed with samtools (0.1.18) (*18*) using only properly paired and uniquely mapped reads. We kept SNPs with variant quality of 30 and genotype quality of 20 and minimum coverage of 10✕, for a total of 60,251 SNPs. Using the concordance procedure described above, we computed concordance rates between the exome dataset and the RNAseq dataset for 30,850 SNPs called in both datasets. The mean concordance rate across individuals is 99.01%. There is one outlier individual with concordance rate of 94.8% (although its concordance rate between RNAseq and genotyping is 99.27%), all other individuals have concordance rate above 98%.

#### 6. Checks in the FCQ dataset

Many additional analyses were performed on the FCQ data to insure the robustness of the results:

- We evaluated differences in diversity between CS and HRR in FCQ (Figure S1), replicating the documented observation of decreased diversity in CS.
- To ensure that the differences in the effects observed in FCQ are not due to biases in the RNAseq data, all results were derived with both RNAseq SNPs, and re-sequencing data of exomes in 96 individuals (Figure S2A, Figure S10). Furthermore, as no significant differences were found with these two samples of different size, the measure of enrichment used to assess the differential mutational burden, is robust to sample size. To evaluate the effect of private variants (Figure 2B, Figure S9), we used the exome sequencing dataset of 96 individuals.
- We evaluated the effect of confounding factors (Table S2-S5). The results are robust to GC content, expression levels and between-species neutral substitution rates (Table S3). Furthermore, we regressed the number of mutations of different types per exon on the recombination rate per exon, controlling for GC content, expression levels, exon size and total SNP density (Table S2). The effect is seen for all mutation types, with no marked differences between transitions and transversions or for mutations towards GC (Table S4), excluding the possibility that GC-biased gene conversion is responsible for the differences seen between recombination environments. More details for controlling for GC content are given below. Finally, we tested the effect using a wide array of recombination rate thresholds used to define CS and HRRs (Table S5).
- We computed OR for non-synonymous and damaging mutations for different frequency classes, to verify that the enrichment of potentially deleterious mutation is not only due to an enrichment of rare variants, that include more non-synonymous variants (Figure S3).

Similar checks were performed in the 1000 Genomes populations. In particular, we performed analyses using the highly covered exons from the RNAseq datasets as well as using all exons where SNPs were called in the 1000 Genomes populations (Figure S2). For other checks (confounding factors, frequency classes) the results obtained are generally the same as in the FCQ, therefore only the FCQ results are shown.

#### 7. Controlling for GC bias

We controlled for variation in GC content in the genome in different ways to verify that this variable is not confounding our analyses. When comparing exons with the same GC content between CS and HRR, the effects remain significant, with the exception of the High GC content class, for non-synonymous (1.07 [0.99;1.15]) and damaging (1.04 [0.92;1.18]). These non-significant results are likely due to the small number of exons in CS with high GC content, and hence the small number of mutations in this class, leading to a lack of power to detect a significant effect.

To make sure that our effects are not influenced by GC-biased gene conversion, a recombination-associated process that favors the fixation of G/C alleles over A/T alleles, we computed the effects independently for all mutations types, and found that the mutations towards G or C showed the same effect as the mutations towards A or T (Table S4).

We further studied the impact of CpG sites. We identified CpG sites by retrieving 3-nucleotide sequences with the central nucleotide being the position of every variant in the FCQ dataset for which the reference allele is a C or a G. Out of 179,005 potential variants, 100,012 were found to be CpG sites. Overall, we see a significant deficit of CpG mutations in CS (OR = 0.62 [0.59;0.64]) reflecting the lower GC content in CS. When excluding all CpG sites from the analyses, the enrichment of putatively deleterious mutations remains significant (Table S4). Similarly, when excluding mutations within CpG islands, the results remain unchanged (Table S4). These additional analyses confirm that GC/CpG content is not responsible for the significant enrichment of putatively deleterious mutations in CS in the human genome.

### Online Supporting Note 3 – Simulations

#### 1. Overview

In the past, selective interference leading to Muller’s ratchet has been mainly investigated through simulation studies. These studies demonstrated the impact of no recombination on the accumulation of deleterious mutations within genomes (*20-24*). However, most results were produced for haploid genomes (but see (*25*)) and were set up to compared regions of free recombination with regions that entirely lack recombination. Here, we performed additional computer simulations (with SLiM (*26*) and sfs_code (*27*)) to describe the expectations under a model of selective interference between negatively selected mutations in diploid genomes, with recombination environments comparable to the ones observed in human autosomes. Both forward-in-time simulation programs gave similar results for all analyses.

#### 2. Recombination environments

We simulated diploid genomes with a distribution of recombination rates similar to the one observed in the empirical data. For 250 individuals (500 haplotypes), we simulated exon-like sequences with non-synonymous mutations, with the mutation rate µ = 2×10^-7^ per base and N_e_ = 1000, chosen to minimize computing time while getting diversity data comparable to human data. We tested models with the overall recombination rate being r = µ or r = µ/2, to evaluate whether the relationship between r and µ had an impact. We defined three recombination environments: coldspots (CS), high recombining regions (HRR) and regions in between. In the human exome, CS and HRR contain 4.1% and 58.6% of recombination events, respectively, according to values in the deCODE map. These values were used in the simulations to match the human genetic map. For each genome, we simulated 75 fragments of 200Kb, with CS and HRR of 95 Kb and 28 Kb, respectively, and 77Kb of regions in between HRR and CS. We also simulated a null model with constant r, to insure that the effects seen do not reflect the difference in region length between CS and HRR. Finally, we also simulated modified CS and HRR, such that their recombination rate matches the African recombination rates within CS and HRR better (Online Supporting Note 1, Figure S11).

#### 3. Models of selection and demography

Our simplest model is a constant population size model with the distribution of fitness effects estimated by Eyre-Walker and colleagues (*28*), using their model without correction for demography (EW model). Simulations were performed with the same distributions of selection coefficients across CS and HRR, with proportion *p* = 75% of variants attributed a selection coefficient from a gamma distribution of mean *s =* -0.0294. We also used other scenarios to model European and African human data, with parameters taken from the EA and AA models from (*29*), a study that inferred both selection and demographic parameters simultaneously. Finally, we simulate data under a model without selection (NTR), a control under selective neutrality but with the human-specific recombination map. The description of these models is shown in Table S6. For each model, we generated 100 replicates. Each replicate took between 15 and 55 hours to run, depending on the model and on the simulation program.

Odds ratios are used to estimate the differential mutational burden between CS and HRR for rare and non-synonymous mutations (Material and Methods). Mutations with derived allele frequency (DAF) below 0.01 are labelled as *rare*. Mutations with s larger than -1/2N_e_ (ns) are effectively neutral, and others are negatively selected (nsneg). To model damaging mutations, we chose the threshold of below s = -0.01 (nsdam), to match the number of non-synonymous damaging variants in the empirical data. Simulated CS have a higher proportion of *rare* and *nsneg* mutations than simulated HRR for all models of selective interference with r = µ. Results for r = µ/2 are highly similar, although the effect is somewhat reduced (Table S7A). The neutral model with varying recombination rates does not show an enrichment of rare neutral variants in CS, indicating that the reduction of N_e_ in low recombination region alone does not account for the excess of rare variants.

#### 4. Models of background selection with and without interference

We simulated various models of background selection, by changing the proportion *p* of non-synonymous variants and the mean selection coefficient *s* from a gamma distribution. We make *p* take value from 0.1 to 0.75 and *s* takes values from -0.0294 to -0.3. We find that the effects on both rare and deleterious diversity increases with *p* (Figure 4). We simulated a scenario with *p* = 0.1 and *s* = -0.3, such that the overall selective pressure acting on the loci is approximately the same as for *p* = 0.75 and *s* = -0.0425 (*28*). Interestingly, this model did not performed better than the scenario with *p* = 0.1 and *s* = -0.0425, suggesting that small proportion of strongly selected mutations is unlikely to cause the difference in mutational burden observed in the data.

Scenarios modelling background selection with no interference further confirm this result: we simulated a single motif of 10 bp in the centre of each region, where all sites have fixed *s* = 0.05, 0.1 or 0.15. These models are designed not to contain linkage between negatively selected sites because the size of the region is too short for many deleterious mutations to occur and interfere with each other. These scenarios do not predict differences in the proportion of rare neutral mutations between CS and HRR (Figure 4), indicating that background selection acting at one independent locus does not lead to an enrichment of rare variants in regions with reduced recombination rate. Therefore, these results show that many mutations with small effects are required to cause the patterns observed, which likely result from reduced efficiency of selection in CS due to interference between these mutations.

#### 5. Simulation of various scenarios to explain population differences

In analyses of the genomic data, differences in the empirical distribution of odds ratios (OR) per individual are observed (Figure 4, Figure S9). They may be related to changes in N_e_, such that smaller populations are even less efficiently purging variants from coldspots than large populations. However, none of the simulated scenarios described above showed significant differences in the distributions of individual OR computed for rare variants. In particular, no significant shift in the OR distribution for rare variants was seen between EA and AA scenarios (Table S6). We also simulated scenarios with population splits, with and without migration between populations and with the dominance coefficient *h* = 0.1. These models also failed to show significant differences between individual OR distributions. To try to understand the observed differences between FCQ and other populations for distributions of individual OR for rare and non-synonymous variants (Figure 4, S9 and S10), several more complex demographic scenarios, with severe bottlenecks and/or with rapid expansions were tested with both SLiM and sfs_code, but none recapitulated the significant difference between distributions seen between the FCQ population and the others (Figure 4). The only scenarios that caused significant differences are when changing the recombination rates within CS and HRR, by increasing the rate in CS and decreasing the rate in HRR (Figure S11). However, the shift in the distribution is very weak and hardly comparable to the differences observed in the empirical data, making it unlikely that the differences in recombination rates alone cause the effect seen between populations. Furthermore, this would be a potential explanation for differences between Africans and non-African populations, but not between Asian, European and French-Canadian populations for which recombination rates are highly similar, therefore differences in recent demographic history is more likely to underlie the differences observed.

One possible explanation for the fact that the simulations do not seem to recapitulate the patterns observed is that the simulation tools fail to adequately model recent population demography of human populations. Although changes in population size can be modelled with these tools, modelling spatial expansion may be essential to get more realistic simulation results. Unfortunately, although spatially explicit simulation tools allowing for complex demography and variation in recombination have been developed (*30*), selection models are not yet included. Furthermore, most simulation tools assume Poisson variance in reproductive success, an assumption often violated in populations with high fertility rates, as it was the case during French-Canadian expansion (*31, 32*). When only few parents contribute to the next generation (*33*), the larger than Poisson variance in family sizes introduces additional stochasticity, causing strong intergenerational genetic drift. Therefore, extensions of simulation methods in the future, to include these additional demographic parameters in flexible simulation frameworks with recombination and selection, will hopefully be informative to understand the impact of complex demography on selective interference. In any case, these results suggest that complex demographic processes, not generally accounted for in population genetics models of human peopling (*34*), may need to be considered to understand the differences in the genomic distribution of deleterious genetic variation.

#### 6. Effect of sample size

We computed our statistics with subsets of individuals simulated to test the effect of sample size on the enrichment statistics. We resampled 125 individuals (half of the number simulated) in 100 replicates and did not find any significant difference in the results obtained (Table S7A).

#### 7. Haplotype load and effect of phasing

We estimated the haplotype load by evaluating the proportion of variants of a given type per haplotype. We counted the number of replicates where the haplotype load is smaller in CS than HRR, to obtain a one-tailed *p*-value (Table S7B). Models with interference are predicted to result in a higher haplotype load in CS for *rare, nsneg* and *nsdam* mutations after correcting for SNP density. However, background selection without interference does not lead to differential haplotype load between recombination environments. We also tested the expectation of the haplotype load under neutrality, to ensure that the differential linkage between CS and HRR is not biasing the expected distribution of haplotypes. These results strongly suggest that the effect observed in the genomic human data is caused by reduced selection efficiency due to interference between mildly deleterious variants.

Because LD-based phasing of rare variants remains technically challenging, we verified that the phasing procedure used in the 1000 Genome data does not influence our results in the haplotype analyses. In particular, the impact of phasing in HRR and CS can be different. We thus verified by simulation that the difference in haplotype load between CS and HRR observed here is not due to potential phasing biases due to low power to phase rare variants. For our 100 NTR simulation replicates, we rephrased the haplotypes using ShapeIt2. We took chunks of same length (25Kb) in simulated CS and HRR and looked at the number of haplotypes with 2 rare mutations and more (MAF < 0.01) in the real haplotypes and the phased haplotypes, N2r and N2p, respectively. There is a phasing bias for both CS and HRR, with the number of haplotypes with 2 mutations and more being reduced by phasing (ie. N2r > N2p), showing that phasing does even out rare mutations across haplotypes (Figure S11B) but no significant difference between CS versus HRR was found in this phasing bias (Figure S11C). These results mean that the effect seen on more loaded haplotypes in CS is not dues to phasing errors, and even suggest that the phasing is probably leading to an underestimation of the increased haplotype load in CS.

### Online Supporting Note 4 – Additional Analyses

#### 1. Variant Functional Annotations

Annotations of synonymous and non-synonymous varinats for the FCQ dataset were obtained from three different sources (as of february 2013): Seattleseq (snp.gs.washington.edu/SeattleSeqAnnotation134), dbSNP (dbSNP135), and wAnnovar (wannovar.usc.edu) (*35*). As these tools consider different databases of transcripts, a SNP is annotated as ‘nonsense’ if at least one annotation tool annotated it as ‘stop gain’ or ‘stop loss’. Similarly, a SNP is annotated as ‘missense’ if at least one annotation tool characterized it as ‘missense’. We note that discordance between these 3 tools is 0.43%. The 1000 Genomes variants were annotated by the Variant Annotation Tool (VAT) and these annotations are used for all results reported. However, the 1000 Genomes variants were also re-annotated using the pipeline used for FCQ variants to ensure results are robust to both annotation pipelines. We find that 2.43% of annotations differ between VAT and the new annotation pipeline, with the vast majority (65.05%) of these discordant variants being annotated as synonymous/intronic with VAT and non-synonymous with the Seattleseq/adSNP/wAnnovar pipeline. We verified that this small fraction of discordant annotated variants is not driving the differences seen between populations for non-synonymous mutations (Figure S12).

The functional impact of non-synonymous mutations was obtained with widely used prediction tools: PolyPhen (*36*) and SIFT(*37*), which both have high sensitivity but low specificity (*38*). We thus used a combination of the two methods to reduce the number of false positives. To estimate the level of constraint at nucleotides, we retrieved GERP conservation scores for all positions within the human exome. GERP scores were obtained from the Sidow lab website (http://mendel.stanford.edu/SidowLab/downloads/gerp/index.html). To reduce false positives, we further obtained PhyloP scores from UCSC Genome Bioinformatics website (http://hgdownload.cse.ucsc.edu/goldenpath/hg19/phyloP46way/). We annotated a mutation as constrained if GERP > 3 and PhyloP > 1. Sites with PhyloP > 1 are the top 10% of conserved sites in the human genome. The GERP threshold was chosen to be comparable to the PhyloP threshold, such that the overall proportion of SNPs and missense SNPs inferred as constrained is the same for both methods. We compared GERP and PhyloP conservations scores with Polyphen and SIFT predictions in the FCQ dataset. Overall, more than half non-synonymous mutations and a large proportion of damaging mutations (78.2%) are conserved according to both conservation scores, although GERP predictions concord slightly better with functional annotations than PhyloP predictions. We thus decided to use GERP predictions to classify exons in conservation categories.

#### 2. Recombination and expression

Expression level of a gene is one of the best predictors of its evolutionary rate (*39*) with the efficiency of selection being weaker in lowly expressed genes. Therefore, we controlled for expression levels in our analyses (Table S2 and S3) and showed that our results are robust to variation in expression levels. However, we note that when analysing highly covered exons, the analyses are biased towards constitutively highly expressed genes. Interestingly, the effects are generally larger for this set of exons relative to all exons in the 1000 Genomes dataset (Figure S2), suggesting that relationship between mutational load, recombination and expression may be of importance. Furthermore, it has been reported that within-gene recombination rates appear to correlate with transcription patterns, such as expression breadth and allelic expression (*40*). We observed a weak but significant negative correlation between recombination rates and mean gene expression in the FCQ data (Table S2), providing further support for a negative association between recombination and transcription in humans.

#### 3. Effect of fixed polymorphism

Variants fixed in a population are excluded from that population, but are kept in the others if they are segregating, and could contribute to the per-individual counts. To verify whether these variants can have an influence in our analyses and could explain population specific differences, we excluded variants where the derived allele is fixed in one population, but is still segregating in the other population from our analyses. This led to 2140 variants from CS and HRR being excluded from the 1000G dataset (0.52%), and 651 from the FCQ dataset (0.24%). We re-computed the per-individual analyses, and found these changes to make no difference in the per-individual distributions (Figure S12). Thus, derived alleles fixed in a population but not in another thus have little impact on the OR distribution per individual.

#### 4. Introns and exon sizes

We computed the distribution of exon and intron lengths in high recombining regions (HRR) and in coldspots (CS). Exons are larger in HRR than in CS and the length distribution are significantly different (Kruskal-Wallis chi-squared = 2089.667, df = 1, p-value < 2. ×10^-16^). The mean length of exons in HRR and CS are 534 bp and 284 bp, respectively. Conversely, introns are larger in CS than in HRR (Kruskal-Wallis chi-squared = 57.495, p = 3.388×10^-14^). Excluding introns longer than 10Kb, the mean length of introns in CS and HRR are 2.313 Kb and 2.260 Kb, respectively (p = 8.303×10^-12^). This confirms the overall negative relationship between intron length and recombination in humans, previously observed on a small set of introns using large-scale recombination rates (*41*). It has been suggested that strong selection favours deletion bias and intron contraction in humans and Drosophila (*42-45*), therefore these results are consistent with the efficiency of selection being reduced in human CS, leading to larger intron sizes in these genomic regions.

We note that exons within a gene can have quite different genomic properties, and variation in recombination rates, conservation levels, GC content and divergence levels between exons is observed. Therefore, conservation and haplotype analyses are done at the exon level in this study.

#### 5. Pseudogenes, Gene Ontology and Disease Analyses

##### Pseudogenes

We retrieved coordinates of the 17,216 regions annotated as pseudogenes from UCSC Tables (http://genome.ucsc.edu/cgi-bin/hgTables), covering 26,222 Kb of sequence. In total, 36% of these regions are in CS and 14% are in HRR. Pseudogenes overlap 9,411 Kb (0.89%) of CS and 3,610Kb of HRR (0.57%). These differences are significant (OR = 1.58 [1.52;1.64]). This result raises the possibility that more genes have lost their protein-coding ability in low recombining regions of the human genome, although an alternative explanation is that more gene duplication were retained within CS throughout evolution, making gene function redundant and not required for survival.

##### Gene Ontology

To further characterize the potential impact on phenotypes, we performed a gene set enrichment analysis, using human gene annotations from the Gene Ontology database and used PANTHER (*47*) and WebGestalt (*48*) resources. For WebGestalt, the reference list is the list of 21,987 genes included in the gtf file used to annotate exons (ftp://ftp.ensembl.org/pub/release71/gtf/homo_sapiens/Homo_sapiens.GRCh37.71.gtf.gz). PANTHER does not accept a reference list larger than 20,000 genes; the default Homo sapiens gene set was therefore used. The submitted gene list contains all annotate CS genes. Classification terms for “Biological Process” and “Molecular function” hierarchy were selected from the Gene Ontology (GO) vocabulary (*46*) if they were found significantly enriched with both tools, with bonferroni correction applied in both. Highly significant GO terms found with WebGestalt but missing from PANTHER GO term list are also reported. CS are enriched for genes involved in essential biological processes such as cell cycle, mitosis, protein metabolic processes, mRNA processing, organelle organisation and microtubule-based processes. Most proteins coded by CS genes are binding proteins (nucleotide, RNA and ATP binding), ligase or transferase (Table S8).

##### Cancer mutations

Somatic mutations from cancer genomes in coding regions were retrieved from the COSMIC database (http://cancer.sanger.ac.uk/cancergenome/projects/cosmic). These mutations where divided into segregating and non-segregating mutations, depending on whether they are found in the 1000 Genomes dataset. Non-segregating cancer mutations from COSMIC are found to be enriched within low recombining regions (Table S9). It should be noted that an important fraction of reported somatic mutations in cancer genomes are ‘passenger’ mutations, and only a subset will contribute to tumour progression. Furthermore, there is substantial ascertainment bias in COSMIC, potentially confounding analyses. For many samples, mutations are reported in candidate genes only, and the result might reflect that candidate genes tend to be enriched in coldspots. This supports results from the Gene Ontology analyses, suggesting that coldspots are likely to be enriched for fundamentally important genes in mitosis, necessary for genomic integrity and stability at the cellular level.

##### Clinically relevant mutations

Clinical variants with potential and validated pathogenicity were retrieved from the curated ClinVar database (*49*) (https://www.ncbi.nlm.nih.gov/clinvar). Variants that are not segregating 1000 Genomes populations (which represent the vast majority of ClinVar variants), or that are found at a MAF lower than 0.01 in these populations, were classified as *rare*. ClinVar variants with MAF > 0.01 were classified as *segregating* variants. ORs were computed by comparing these rare variants to non-disease variants with MAF in 1000 Genomes populations lower than 0.01 between CS and HRR (Figure 6). We find that rare clinvar variants are in excess in CS after correcting for MAF, suggesting that decreased efficiency of selection gives rise to an excess of disease causing mutations in low recombining regions. Conversely, a lack of variants with MAF > 0.01 implicated in human disease is notable in CS. These common variants were likely identified by genome-wide association studies (GWAS). This result may reflect a bias in GWAS towards higher power of discovery of common variants in HRR. The result for rare variants were replicated using the Humsavar database (http://www.uniprot.org/docs/humsavar), which are disease mutations identified from the Universal Protein Resource (Table S9). However, the result for variants with MAF > 0.01 is not seen in Humsavar. Furthermore, sensitive and ultra-sensitive motifs (*50*) are enriched in CS compared to HRR. These motifs have been previously examined for the presence of inherited disease-causing mutations from HGMD (Human Gene Mutation Database) and a ∼40- and ∼400- fold enrichment of disease-causing mutations in sensitive and ultrasensitive regions was found, respectively.

##### GWAS hits and genotyping chips

The catalog of published GWAS hits was retrieved from NHGRI (http://www.genome.gov/gwastudies/). We compared the number of hits found in CS and HRR, correcting by total diversity (from the 1000 Genome whole genome SNP data) and found that GWAS hits are highly under-represented. Although CS are regions of extended LD where tagging should be more efficient, the tagging SNPs yielding high power in association studies are in general common SNPs that are in strong LD with the causal SNP. Therefore, in low recombination regions, most variants at lower frequencies are likely to be poorly tagged by common markers on genotyping chips used for performing GWAS. To test this hypothesis, we retrieved the list of markers from three genotyping SNPs, highly used for GWAS: Affymetrix SNP Array 6.0, Illumina Omni 1M and Illumina Omni 2.5M. All chips are enriched for tagged markers in HRR, even when looking at common variants (>10%) only. After controlling for this bias towards HRR in genotyping chips, there is no significant difference in the distribution of GWAS hits between HRR and CS is seen (OR = 1.05 [0.99;1.11]). Therefore, it is likely that the bias in GWAS hits is partially caused by the bias in distribution of markers in genotyping chips, although it is expected that the effectiveness of tagging is higher in CS, explaining why the design of genotyping chips is biased towards high recombining regions. Furthermore, it has been showed that ascertainment bias will likely erode the power of tests of association between SNPs and complex disorders, and that this will affect the power to detect associations when the variants that cause the inflated risk are rare (*51*). These results thus suggest altogether that the common variant model on which GWAS using commonly used genotyping chips has been based is not appropriate to find most potential disease variants in CS and that a substantial proportion of undiscovered mutations associated with disease phenotype may be segregating in conserved low recombining regions. This is a topic that goes beyond the scope of this article, but this is an important problem worthy of additional study.

## Supplementary Figures

**Figure S1.**
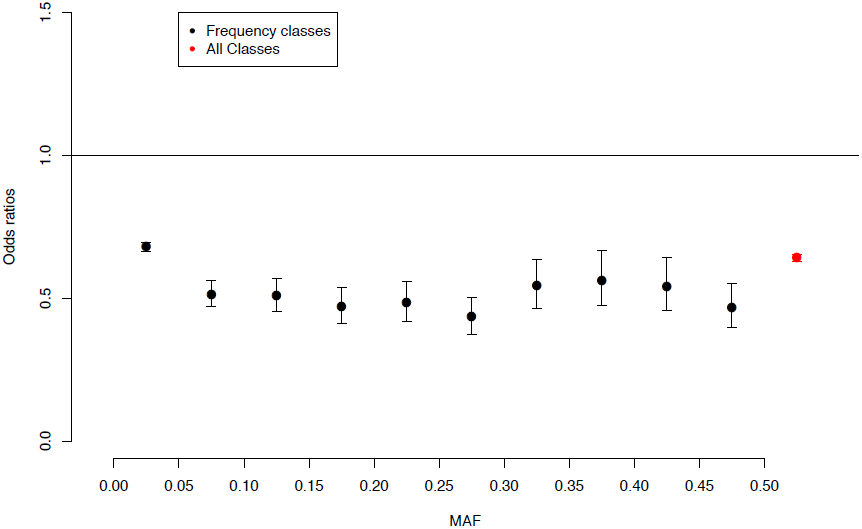
Comparison of levels of diversity between Coldspots (CS) and High Recombining Regions (HRR) for SNPs in the FCQ dataset. Odds ratios (OR) are computed to compare SNP density between CS and HRRs for all SNPs (red) and SNPs divided in different allele frequencies classes (black). OR < 1 means that diversity is greater in HRR than in CS. We confirm the lack of diversity in CS relative to HRR, in line with previous evidence that diversity is reduced in low recombining regions due to background selection. The effect is seen for all frequency classes and does not differ significantly between classes of SNPs with MAF > 0.05. The class of variants with MAF < 0.05 shows a slightly smaller effect than the other frequency classes.

**Figure S2.**
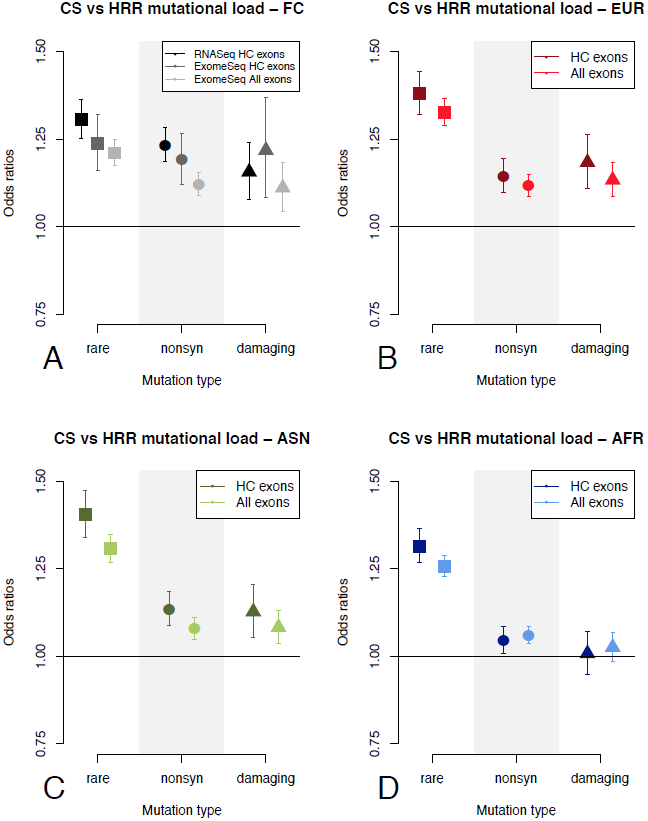
Differential mutational burden in genomic subset of the data. Differential burden is computed using odds ratios (OR), representing the relative enrichment of a category of variants compared to all variants in CS vs. HRR for (A) RNA and exome sequencing of French-Canadians and for exome sequencing of (B) Europeans, (C) Asians and (D) Africans, from the 1000 Genome Project. Variants are categorized as rare (MAF < 0.01 in a population), non-synonymous (missense and nonsense) and damaging (predicted by both SIFT and Polyphen2). Highly covered exons (HC exons) have coverage above 20X for each position within the exons in all datasets. The set of exons analysed does not affect the results and the exome dataset in French-Canadians replicates the results found in RNAseq.

**Figure S3.**
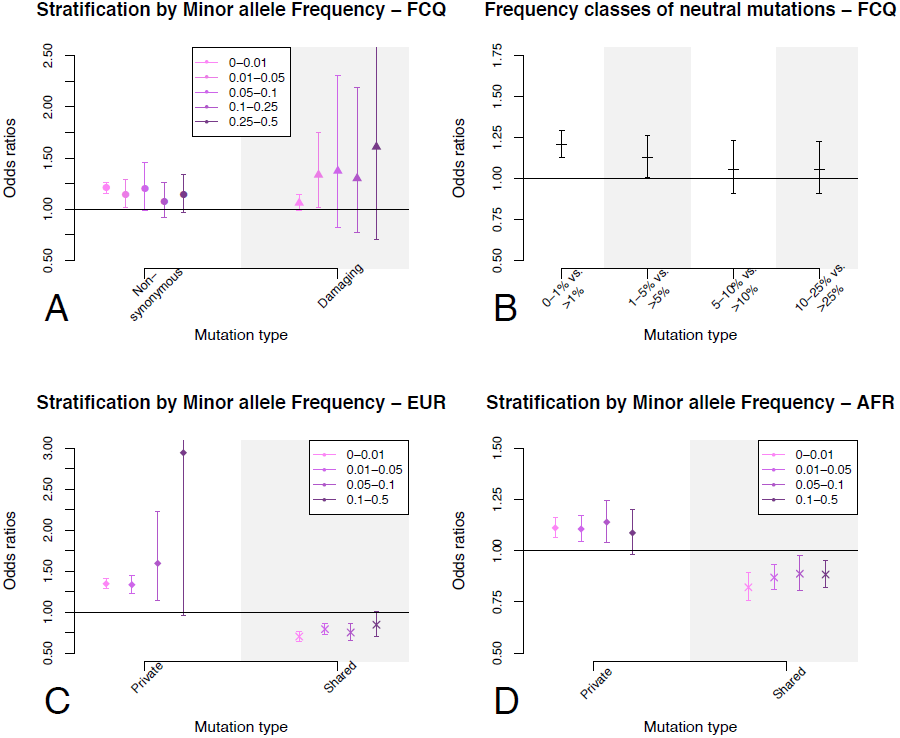
Minor allele frequencies (MAF) impact on odds ratios between CS and HRR. Impact of MAF on the effects for functional mutations in the FCQ RNAseq dataset (A,B) and for private and shared variants in EUR (C) and AFR (D). (A) The enrichment of non-synonymous and damaging mutations in CS remains significant for MAF < 0.05, indicating that the excess of rare variants in CS does not drive the effect for non-synonymous and damaging variants. (B) Neutral variants with MAF < 0.05 are enriched in CS compared to more frequent variants, indicating that neutral diversity contributes to the excess of rare variants in CS. (C,D) The enrichment of private mutations in CS and of shared mutations in HRR remains significant for MAF < 0.1 in both EUR and AFR, indicating that these effects are not driven only by differences of allele frequencies between shared and private variants.

**Figure S4.**
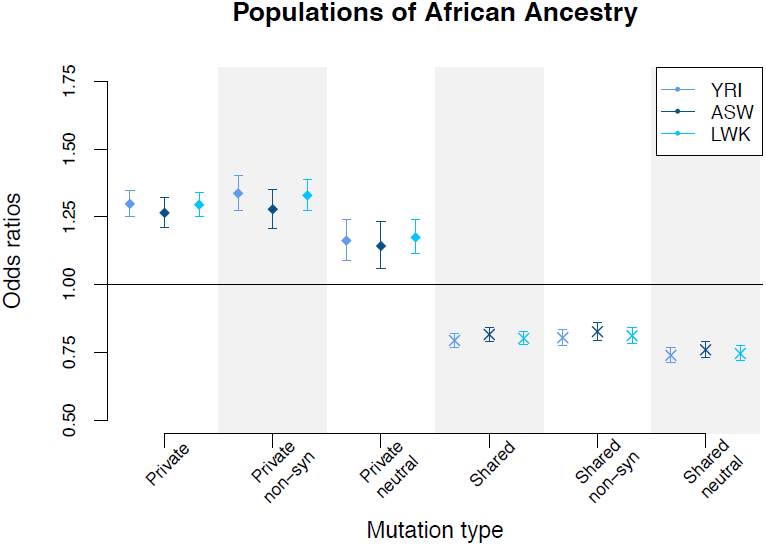
Effects for private and shared variants between African sub-populations. Comparison of closely related populations of African ancestry. ORs are computed based on private and shared variants called in 88 Yoruba in Ibadan from Nigeria (YRI), 97 Luhya in Webuye from Kenya (LWK), and 61 Americans of African Ancestry (ASW).

**Figure S5.**
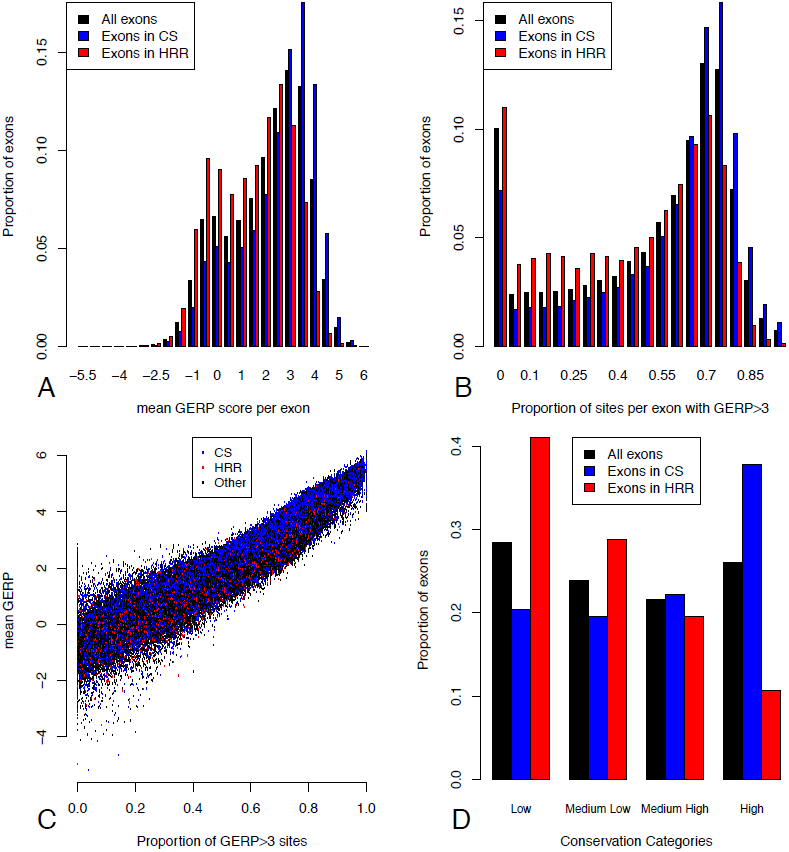
Distribution of conservation across exons measured by GERP scores. Mean GERP score per exon (B) Proportion of constrained positions (GERP > 3) per exon (C) Scatter plot of mean GERP by the proportion of constrained positions for all exons (D) For each measure of conservation per exon, exons were grouped into 4 categories of equal sizes. Only exons that were concordant between the two classifications were kept in analyses within conservation categories, to minimize the effect of outliers with one of the two measures. Characteristics of exons in these four conservation categories in terms of average GERP score per base pair (bp) and number of constrained sites per bp (GERP > 3) are reported in Figure 3C.

**Figure S6.**
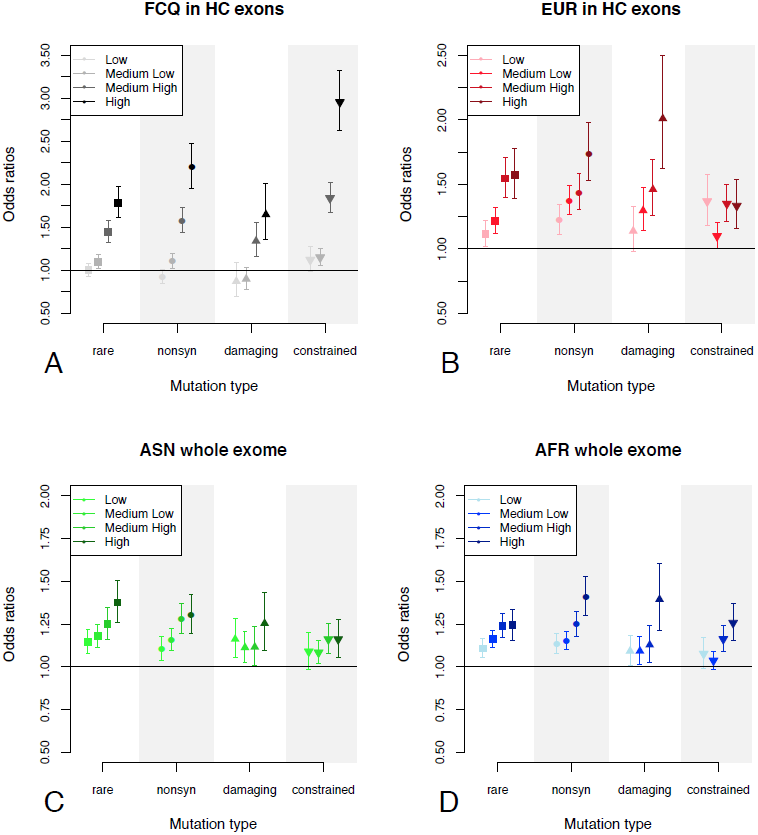
Differential Mutational Burden in Conservation categories. Differential mutational burden between coldspots (CS) and highly recombining regions (HRR) for rare (MAF < 0.01), non-synonymous (nonsyn), damaging and constrained variants in (A) French-Canadians and (B) Europeans for Highly Covered (HC) exons, and in (C) Asians and (D) African for the whole exome. Results for Europeans in the whole exome are presented in Figure 3A. Results for Asians and Africans in HC exons (not shown), are similar to European results. For all populations and exon datasets, the Medium High and High conservation categories always show a significant enrichment for potentially deleterious mutations in CS.

**Figure S7.**
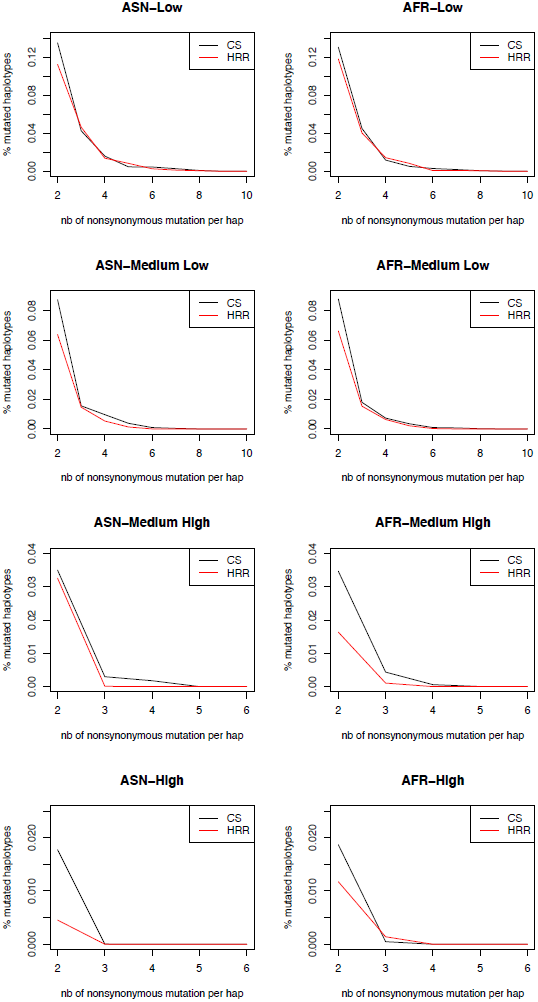
Haplotype load of non-synonymous variants in CS and HRR in Asians and Africans in the different conservation categories. Haplotype load is computed as described in Material and Methods. Haplotype load in Europeans in presented in Figure 3B, with characteristics of conservation categories shown in Figure 3C.

**Figure S8.**
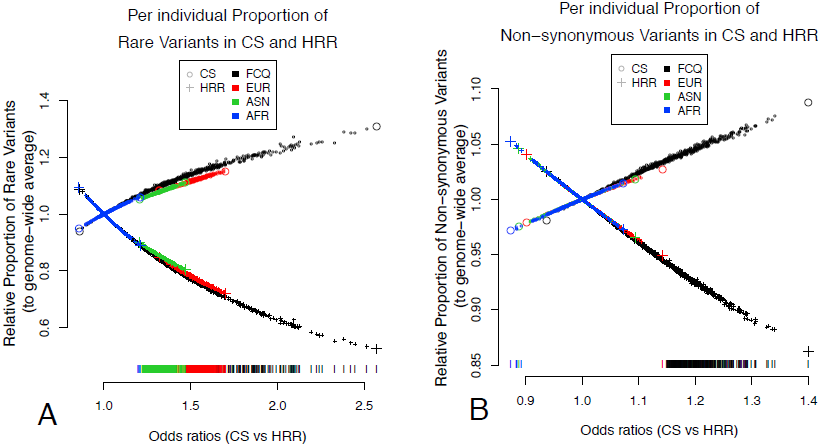
Per individual differential mutational burden across populations. Comparison of proportions of (A) rare and (B) non-synonymous mutations between coldspots (CS) and high recombining regions (HRR). For each individual (ordered by their OR values), the relative proportions of rare or non-synonymous mutations in CS and HRR is shown, computed by dividing CS and HRR proportions by genome-wide proportions of rare or non-synonymous variants within each individual, to adjust for differences across individuals. The larger symbols represent individuals with the minimum and maximum OR values in each population. Ticks at the bottom of the plots show individual OR values significantly different from 1 (two-tailed p < 0.05). French-Canadian data used is the RNAseq dataset (Online Supporting note 2), replication with exome data of 96 French-Canadians is presented Figure S10.

**Figure S9.**
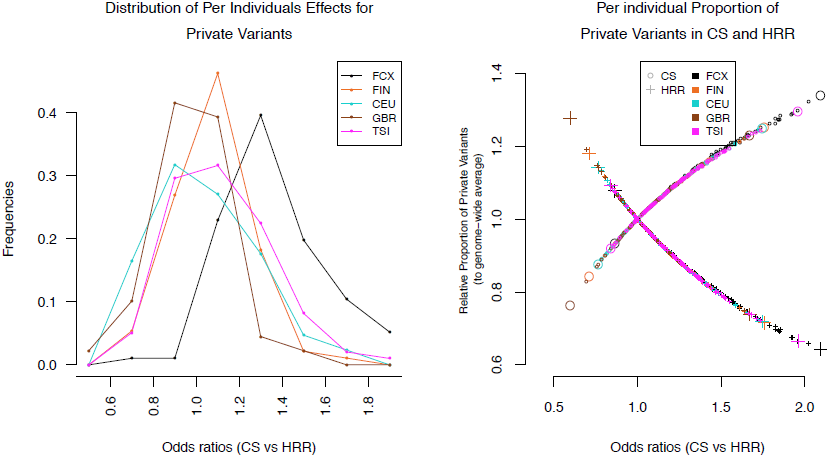
Per individual differential mutational burden across European populations for private variants. Distribution of odds ratios (OR) per individual comparing proportions of private variants between CS and HRR in closely related populations of West European ancestry. OR are computed based on private variants called in the exome sequencing dataset of 96 French-Canadians (FCX), 89 British individuals (GBR), 93 Finns (FIN), 98 Italians from Tuscany (TSI) and 85 European Americans (CEU). The left panel shows the frequencies of individual OR in each population. The right panel shows, for each individual (ordered by their OR values), the relative proportions of private mutations in CS and HRR, computed by dividing CS and HRR proportions by genome-wide proportions of private variants within each individual, to adjust for differences across individuals.

**Figure S10.**
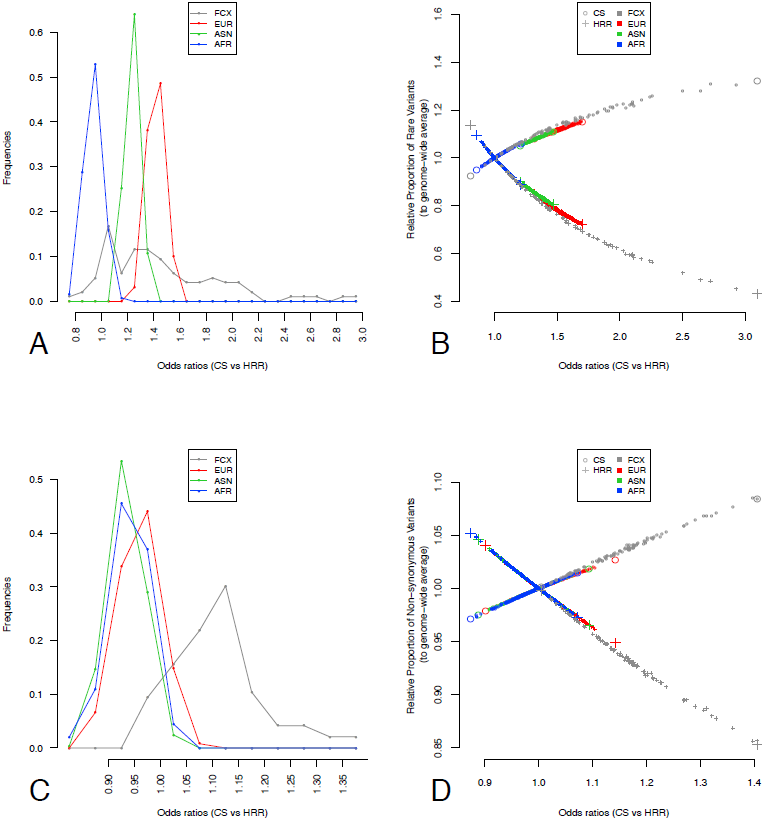
Per individual differential mutational burden across populations with FCQ exome sequencing data. Distribution of odds ratios (OR) per individual comparing proportions of rare (A, B) and non-synonymous (C, D) mutations between CS and HRR. For Europeans, Asians and Africans, the results are the same as shown in Figure 4, whereas French-Canadian results are computed using the exome sequencing dataset from 96 individuals. Further descriptions of the plots are found in Figure 4.

**Figure S11.**
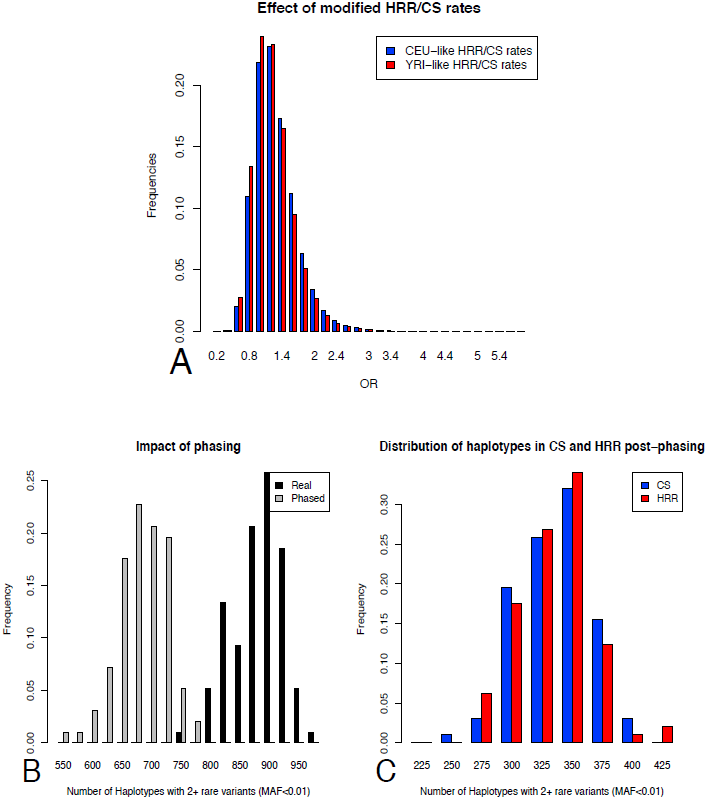
Additional simulations testing the effect of recombination rates and phasing. (A) Distribution of initial and modified CS and HRR, with CS/HRR recombination rate matching the rates in the CEU and YRI maps, respectively (Online Supporting Note 1.4). The distributions are significantly different, but the shift in mean is very weak, and unlikely to cause the large differences we observe between populations in Figure 5. (B,C) Effect of phasing on the distribution of the number of haplotypes with 2 rare mutations and more (MAF < 0.01) in the real haplotypes and the phased haplotypes on chunks of same length (25Kb) in simulated CS and HRR. (B) The number of haplotypes with 2 mutations is reduced by statistical phasing with ShapeIt2 but (C) no significant difference between CS versus HRR was found in this phasing bias.

**Figure S12.**
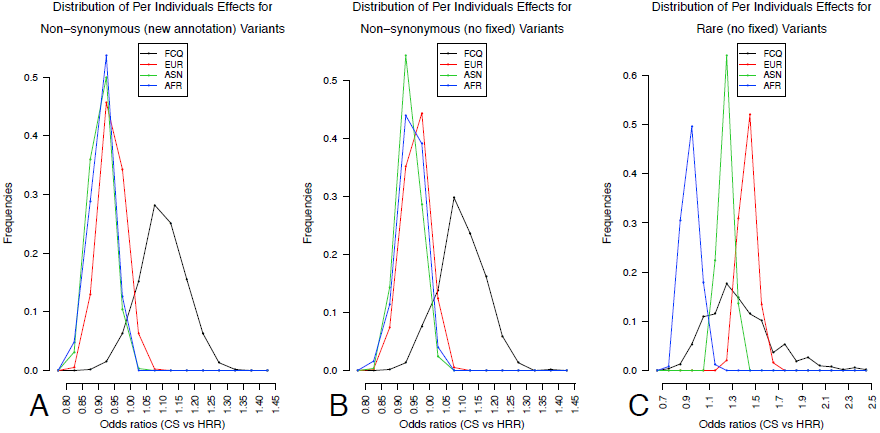
Quality checks on per individual differential mutational burden across populations. Distribution of odds ratios (OR) per individual in French-Canadians, Europeans, Asians and Africans, comparing proportions of (A) non-synonymous variants after modifying annotations in the 1000 Genomes Populations (see Online Supporting Note 4.1) and (B) non-synonymous and (C) rare variants, after excluding mutations that are fixed in one population but still segregating in others, between coldspots (CS) and high recombining regions (HRR). The differences between populations observed in Figure 5 remain the same after correcting for these possible technical differences.

**Figure S13.**
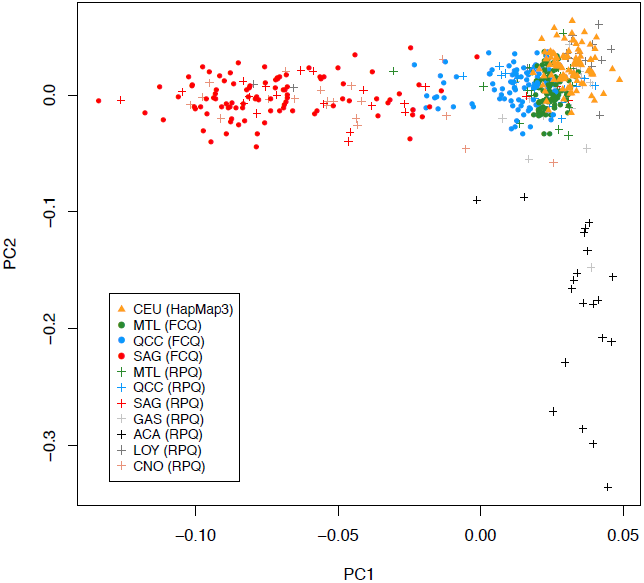
Population structure in regional populations of Quebec. Sampling from the CARTaGENE Project includes individuals from the Montreal area (MTL). Quebec City (QCC) and the Saguenay region (SAG). Regional origin of individuals was confirmed by a principal component analysis of genetic diversity in FCQ individuals compared with genetic diversity within the Reference Panel of Quebec (RPQ) and in the CEU population from HapMap3. Other populations included in the RPQ are : GAS : Gaspesia Region. ACA : Acadians. LOY : Loyalists. CNO : North Shore Region (*52*).

## Supplementary Tables

**Table S1.** Distribution of sequence (A) and SNPs (B) in Coldspots (CS) and High Recombination Regions (HRR) genome-wide and in highly covered (HC) exons.

| Regions | CS (bp) | HRR (bp) |
| --- | --- | --- |
| Whole genome | 1.048.937.114 | 634.243.758 |
| Whole exome | 25.302.008 | 17.768.017 |
| HC exons | 6.906.137 | 2.036.963 |

**B**
| POP<sup>a</sup> | Number of individuals | Number of SNPs<br>Whole Datasets |  |  | Number of SNPs<br>HC exons<sup>b</sup> |  |  |
| --- | --- | --- | --- | --- | --- | --- | --- |
|  |  | TOTAL | HRR | CS | TOTAL | HRR | CS |
| FCQ | 521 | 178,394 | 37,076 | 69,295 | 73,627 | 14,899 | 30,734 |
| EUR | 379 | 142,296 | 34,248 | 48,665 | 69,672 | 12,489 | 29,827 |
| ASN | 286 | 128,697 | 31,372 | 44,292 | 63,726 | 11,358 | 27,562 |
| AFR | 246 | 186,549 | 45,816 | 62,944 | 89,789 | 16,464 | 38,157 |

<sup>a</sup> French-Canadians (FCQ) Europeans (EUR). Asians (ASN) and Africans (AFR).

**C**
| POP<sup>a</sup> | Number of individuals | Number of non-synonymous SNPs<br>HC exons |  |  | Number of rare SNPs<br>HC exons |  |  |
| --- | --- | --- | --- | --- | --- | --- | --- |
|  |  | TOTAL | HRR | CS | TOTAL | HRR | CS |
| FCQ | 521 | 31,472 | 6,488 | 14,984 | 42,627 | 10,097 | 22,530 |
| EUR | 379 | 37,712 | 6,536 | 16,614 | 48,045 | 8,111 | 21,447 |
| ASN | 286 | 34,322 | 5,949 | 15,307 | 44,676 | 7,511 | 20,196 |
| AFR | 246 | 44,040 | 8,065 | 19,117 | 51,863 | 8,871 | 23,111 |

<sup>a</sup> French-Canadians (FCQ) Europeans (EUR). Asians (ASN) and Africans (AFR).

**Table S2.** Summary of linear regression models (FCQ dataset)

| Variable | Rec rates<sup>a</sup> | GC content | Expression | Exon Size | SNPs/Kb | R<sup>2</sup> |
| --- | --- | --- | --- | --- | --- | --- |
| **SNP/Kb** | + *** | + *** | + *** | NS | NA | 0.038 |
| **Average MAF** | + *** | + *** | + *** | + *** | NA | 0.046 |
| **Expression<sup>b</sup>** | - *** | + *** | NA | NS | NA | 0.016 |
| **Density (SNP/Kb) for** |  |  |  |  |  |  |
| Constrained | - *** | NS | NS | - *** | + *** | 0.267 |
| Non-synonymous | - * | - ** | - *** | - *** | + *** | 0.538 |
| Damaging | NS | NS | - *** | - *** | + *** | 0.151 |
| Private | - *** | + *** | NS | NS | + *** | 0.387 |

\*\*\* p<0.001; \*\* p<0.01; \* p<0.05; NS : non-significant; +/- = positive/negative correlation; NA : non-applicable.
<sup>a</sup> Average recombination rates per exon in cM/Mb are computed based on the FCQ genetic map.
<sup>b</sup> This correlation is evaluated considering all exons with minimum coverage of 20x in at 50% individuals. It is therefore biased against low expression genes.

**Table S3.**
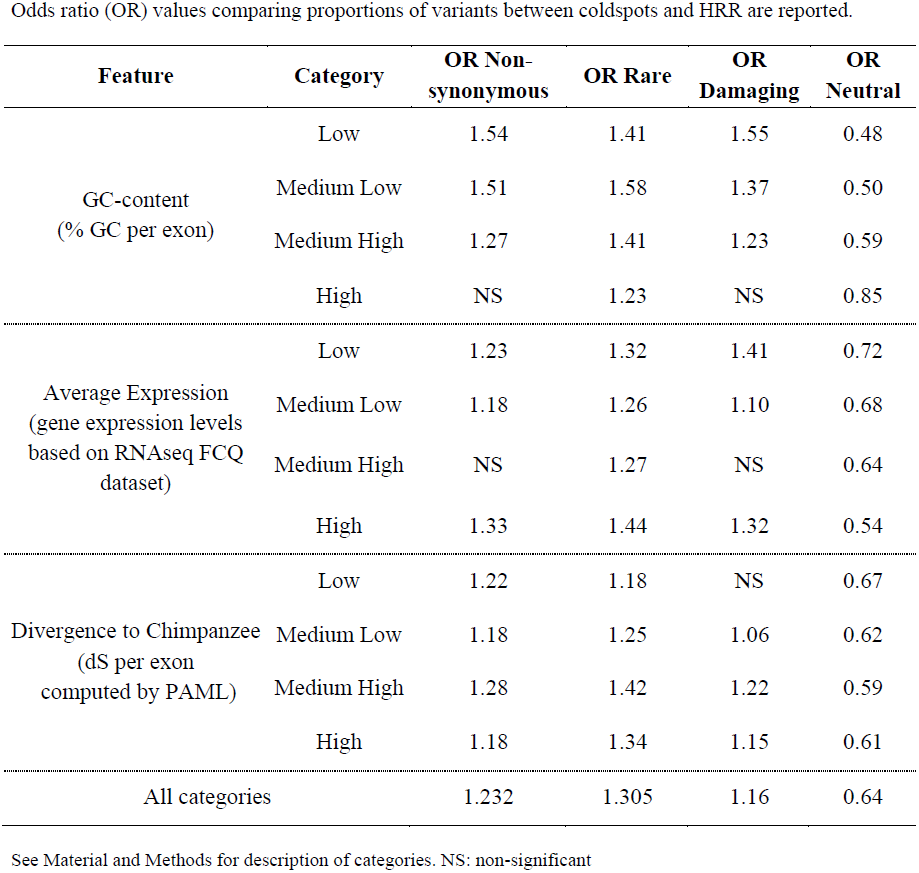
Robustness of the effect to GC-content. gene expression levels and divergence (FCQ dataset)

**Table S4.** Effects for different mutation types (FCQ dataset)

| **Mutation type** | **OR Rare** |  | **OR Non-syn** |  | **OR damaging** |  | **OR neutral [CI** |  |
| --- | --- | --- | --- | --- | --- | --- | --- | --- |
|  | **[CI 95%]** |  | **[CI 95%]** |  | **[CI 95%]** |  | **95%]** |  |
| G→C/C→G | 1.44 | [1.24;1.67] | 1.49 | [1.3;1.71] | 1.44 | [1.14;1.80] | 0.54 | [0.44;0.67] |
| A→G/ T→C | 1.22 | [1.16;1.30] | 1.26 | [1.19;1.33] | 1.24 | [1.13;1.37] | 0.69 | [0.64;0.75] |
| A→C/ T→G | 1.42 | [1.21;1.69] | 1.21 | [1.04;1.41] | 1.27 | [0.99;1.63] | 0.59 | [0.46;0.76] |
| G→A/ C→T | 1.49 | [1.36;1.64] | 1.06 | [0.97;1.16] | 0.88 | [0.74;1.05] | 0.53 | [0.48;0.60] |
| G→T/ C→A | 1.53 | [1.23;1.89] | 1.40 | [1.15;1.71] | 1.37 | [0.95;1.95] | 0.39 | [0.30;0.53] |
| A→T/T→A | 1.25 | [1.18;1.32] | 1.25 | [1.19;1.32] | 1.25 | [1.14;1.36] | 0.68 | [1.63;1.74] |
| Exclusion of CpGs sites | 1.47 | [1.39;1.56] | 1.28 | [1.22;1.35] | 1.20 | [1.09;1.33] | 0.56 | [0.52;0.60] |
| Exclusion of CpG islands | 1.33 | [1.27;1.39] | 1.25 | [1.20;1.30] | 1.15 | [1.07;1.24] | 0.62 | [0.58;0.66] |

**Table S5.** Robustness of the effect to the choice of recombination parameters (FCQ dataset)

| Rec. rates<br>(cM/Mb) |  | Odds ratios CS vs HRRs |  |  | Number of SNPs |  |  |
| --- | --- | --- | --- | --- | --- | --- | --- |
| L | H | Non-synonymous | Rare | Private | CS | HRR | In between |
| 0.1 |  | 1.24 | 1.27 | 1.21 | 9263 | 64367 | - |
|  | 5 | 1.29 | 1.34 | 1.27 | 9263 | 15208 | 49159 |
|  | 10 | 1.25 | 1.33 | 1.26 | 9263 | 10406 | 53961 |
|  | 20 | 1.2 | 1.26 | 1.17 | 9263 | 7517 | 56850 |
| 0.25 |  | 1.19 | 1.25 | 1.2 | 20482 | 53148 | - |
|  | 5 | 1.24 | 1.32 | 1.25 | 20482 | 15054 | 38094 |
|  | 10 | 1.21 | 1.31 | 1.24 | 20482 | 10210 | 42938 |
|  | 20 | 1.16 | 1.24 | 1.21 | 20482 | 7272 | 45876 |
| 0.5 |  | 1.18 | 1.25 | 1.18 | 30789 | 42841 | - |
|  | 5 | **1.23** | **1.31** | **1.26** | **30789** | **14902** | **27939** |
|  | 10 | 1.21 | 1.31 | 1.25 | 30789 | 9903 | 32938 |
|  | 20 | 1.17 | 1.25 | 1.17 | 30789 | 6857 | 35984 |
| 1 |  | 1.17 | 1.22 | 1.15 | 41034 | 32596 | - |
|  | 5 | 1.19 | 1.27 | 1.22 | 41034 | 14770 | 17826 |
|  | 10 | 1.17 | 1.27 | 1.23 | 41034 | 9620 | 22976 |
|  | 20 | 1.13 | 1.2 | 1.13 | 41034 | 6352 | 26244 |
| 2 |  | 1.14 | 1.22 | 1.14 | 49583 | 24047 | - |
|  | 5 | 1.15 | 1.24 | 1.19 | 49583 | 14915 | 9132 |
|  | 10 | 1.15 | 1.25 | 1.21 | 49583 | 9389 | 14658 |
|  | 20 | 1.12 | 1.19 | 1.12 | 49583 | 5892 | 18155 |

Coldspots Regions: SNPs within a 50Kb region with no recombination rate higher than L
High Recombination Regions (HRRs): SNPs within 50Kb of at least two hotspots with rate higher than H
Parameters used in this study (red) were chosen to maximize the overall number of SNPs included in the analyses while minimizing the difference between the number of SNPs in coldspots and in HRRs.

**Table S6.** Demographic and selection models used in simulations.

| Model | Parameters |  |  |  |  |
| --- | --- | --- | --- | --- | --- |
| | $\mu$ | $r$ | Genetic Map | Demography | Mean $s^a$ |
| EW $\mu=r$ | $2 \times 10^{-7}$ | $2 \times 10^{-7}$ | CS/HRR | constant size | -0.0425 |
| EW $\mu=2r$ | $2 \times 10^{-7}$ | $10^{-7}$ | CS/HRR | constant size | -0.0425 |
| AA $\mu=r$ | $2 \times 10^{-7}$ | $2 \times 10^{-7}$ | CS/HRR | expansion | -0.0294 |
| AA $\mu=2r$ | $2 \times 10^{-7}$ | $10^{-7}$ | CS/HRR | expansion | -0.0294 |
| EA $\mu=r$ | $2 \times 10^{-7}$ | $2 \times 10^{-7}$ | CS/HRR | bottleneck + expansion | -0.03 |
| EA $\mu=2r$ | $2 \times 10^{-7}$ | $10^{-7}$ | CS/HRR | bottleneck + expansion | -0.03 |
| EW constant $r$ | $2 \times 10^{-7}$ | $2 \times 10^{-7}$ | constant rate | constant size | -0.0425 |
| NTR | $2 \times 10^{-7}$ | $2 \times 10^{-7}$ | CS/HRR | constant size | 0 |

<sup>a</sup> Negative selection is modelled by a gamma distribution of mean $s$ , with $p=75\%$ of mutations attributed a non-zero selective coefficients
EW : DFE from Eyre-Walker et al. 2006
EA : DFE and demographic model from Boyko et al. 2008 for European-Americans
AA : DFE and demographic model from Boyko et al. 2008 for African-Americans
NTR : No selection

**Table S7.** Differential Mutational and Haplotype Load in CS versus HRR in Simulations.

| Odds Ratios for Different Models (Sample size n=500) |  |  |  |  |  |  |  |  |
| --- | --- | --- | --- | --- | --- | --- | --- | --- |
| Mut Type | EW $\mu=r$ [CI 95%] | | EW $\mu=2r$ [CI 95%] | | constant $\rho$ [CI 95%] | | NTR [CI 95%] | |
| rare | **1.22** | [1.12;1.29] | **1.14** | [1.08;1.2] | 0.97 | [0.9;1.05] | 0.99 | [0.92;1.07] |
| ns | 1.05 | [0.999;1.12] | 1.04 | [0.99;1.09] | 0.99 | [0.94;1.05] | - | - |
| nsneg | **1.1** | [1.04;1.16] | **1.06** | [1.01;1.12] | 0.99 | [0.93;1.06] | - | - |
| nsdam | **1.12** | [1.03;1.19] | **1.08** | [1.02;1.16] | 0.98 | [0.9;1.09] | - | - |
| | AA $\mu=r$ [CI 95%] | | AA $\mu=2r$ [CI 95%] | | EA $\mu=r$ [CI 95%] | | EA $\mu=2r$ [CI 95%] | |
| rare | **1.18** | [1.14;1.23] | **1.14** | [1.09;1.19] | **1.21** | [1.14;1.27] | **1.14** | [1.09;1.22] |
| ns | 1.03 | [0.998;1.07] | 1.02 | [0.98;1.06] | 1.03 | [0.99;1.07] | 1.02 | [0.98;1.05] |
| nsneg | **1.06** | [1.02;1.1] | **1.04** | [1.002;1.09] | **1.05** | [1.01;1.09] | 1.03 | [0.99;1.08] |
| nsdam | **1.09** | [1.01;1.17] | 1.07 | [0.99;1.14] | 1.06 | [0.998;1.14] | 1.04 | [0.99;1.11] |

| (Sample size n=200) | EW $\mu=r$ [CI 95%] | | AA $\mu=r$ [CI 95%] | | EA $\mu=r$ [CI 95%] | |
| --- | --- | --- | --- | --- | --- | --- |
| rare | **1.20** | [1.06;1.26] | **1.15** | [1.07;1.25] | **1.18** | [1.1;1.32] |
| ns | 1.07 | [0.95;1.15] | 1.02 | [0.95;1.08] | 1.01 | [0.97;1.09] |
| nsneg | **1.11** | [1.05;1.2] | **1.05** | [1.01;1.09] | **1.07** | [1.01;1.011] |
| nsdam | **1.12** | [1.02;1.25] | 1.07 | [0.98;1.16] | 1.07 | [1.004;1.13] |

**B**
| Models | Mutation Type |  |  |  |
| --- | --- | --- | --- | --- |
|  | rare | ns | nsneg | nsdam |
| EW $\mu=r$, $p=0.75$ | **<0.01** | 0.13 | **<0.01** | **0.01** |
| AA $\mu=r$ | **<0.01** | 0.11 | **0.01** | **<0.01** |
| EA $\mu=r$ | **<0.01** | 0.05 | **0.01** | **<0.01** |
| EW $\mu=r$, $p=0.1$ | **0.03** | 0.42 | 0.35 | 0.28 |
| EW $\mu=r$, $p=0.5$ | **<0.01** | 0.09 | **<0.01** | **0.01** |
| 10bp deleterious motif ($s = 0.1$) | 0.55 | - | - | - |
| EW constant $r$ | 0.74 | 0.54 | 0.68 | 0.69 |
| NTR | 0.56 | - | - | - |
| NTR rephrased with ShapeIt2 | 0.63 | - | - | - |

Proportion of simulated replicates where haplotypes in HRR have a higher proportion of a given mutation type. Values in bold represent significant results (one-tailed p-value <0.05).

**Table S8.** Gene Ontology Analysis.

| **Terms** | **Number of**<br>**genes<sup>a</sup>** | **WebGestalt**<br>**p-value** | **PANTHER**<br>**p-value** |
| --- | --- | --- | --- |
| **Biological processes** |  |  |  |
| cell cycle | 750 | $1.93 \times 10^{-15}$ | $2.18 \times 10^{-11}$ |
| mitosis | 420 | $1.93 \times 10^{-7}$ | $1.13 \times 10^{-8}$ |
| protein metabolic processes | 1578 | $8.32 \times 10^{-12}$ | $2.50 \times 10^{-9}$ |
| mRNA processing | 246 | $1.22 \times 10^{-3}$ | $7.95 \times 10^{-8}$ |
| organelle organisation | 1065 | $3.56 \times 10^{-14}$ | term not included |
| microtubule-based processes | 246 | $3.08 \times 10^{-9}$ | term not included |
| **Molecular function** |  |  |  |
| Binding | 5154 | $8.64 \times 10^{-8}$ | $5.87 \times 10^{-13}$ |
| nucleotide binding | 1192 | $3.00 \times 10^{-13}$ | $1.96 \times 10^{-14}$ |
| RNA binding | 435 | $2.08 \times 10^{-6}$ | $1.24 \times 10^{-7}$ |
| ATP binding | 772 | $2.77 \times 10^{-13}$ | term not included |
| Catalytic Activity | 2472 | $5.06 \times 10^{-11}$ | $2.75 \times 10^{-26}$ |
| ligase | 279 | $1.54 \times 10^{-9}$ | $1.38 \times 10^{-11}$ |
| transferase | 840 | $5.18 \times 10^{-5}$ | $1.99 \times 10^{-7}$ |

<sup>a</sup>The number of genes is the maximum number reported between WebGestalt and PANTHER

**Table S9.** Overrepresentation of clinically relevant mutations, cancer variants and sensitive genomic regions in coldspots.

|  | COUNTS |  | OR for CS vs HRR |
| --- | --- | --- | --- |
|  | CS | HRR |  |
| Genome-wide SNPs (1000G) |  |  |  |
| rare (<0.01) | 7,445,567 | 5,435,142 | correcting for genome-wide diversity |
| segregating (>0.01) | 4,883,516 | 4,160,183 |  |
| clinvar mutations | 18,103 | 11,961 | 1.18 [1.15;1.21] |
| segregating | 989 | 928 | 0.91 [0.83;0.99] |
| non-segregating and rare | 17,114 | 11,033 | 1.13 [1.10;1.16] |
| humsavar mutations | 11,080 | 8,258 | 1.04 [1.01;1.07] |
| segregating | 7,763 | 6,119 | 1.08 [1.04;1.11] |
| non-segregating and rare | 3,317 | 2,139 | 1.13 [1.07; 1.20] |
| whole exome (pb) | 25,302,008 | 17,768,017 | correcting for sequence length |
| cosmic mutations (point mut.) | 574,072 | 316,080 | 1.275 [1.270;1.281] |
| segregating | 16,254 | 12,648 | 0.902 [0.881;0.923] |
| non-segregating | 557,818 | 303,432 | 1.290 [1.285;1.296] |
| motifs in Khurana et al. (pb) |  |  |  |
| sensitive | 2,087,033 | 963,237 | 1.522 [1.518; 1.525] |
| ultra-sensitive | 165,453 | 17,859 | 6.506 [6.406; 6.607] |
| Genome-wide SNPs (1000G) | 12,329,083 | 9,595,325 | correcting for total number of SNPs |
| GWAS hits | 3,470 | 4,836 | 0.558 [0.535; 0.583] |
| Affy 6.0 Chip | 223,836 | 288,879 | 0.603 [0.599; 0.606] |
| Illumina 1M | 301,582 | 306,208 | 0.776 [0.771;0.780] |
| Illumina 2.5M | 573,726 | 815,220 | 0.548 [0.546; 0.550] |
| all Chips | 869,577 | 1,037,827 | 0.652 [0.650;0.654] |
| Common SNPs (1000G, >10%) | 1,653,476 | 1,566,782 | correcting for number of common SNPs |
| GWAS hits | 2,367 | 3,315 | 0.6756 [0.641; 0.713] |
| Affy 6.0 Chip | 162,275 | 219,378 | 0.701 [0.696; 0.706] |
| Illumina 1M | 203,087 | 248,019 | 0.776 [0.771; 0.781] |
| Illumina 2.5M | 220,439 | 460,997 | 0.450 [0.447;0.453] |
| all Chips | 415,062 | 613,188 | 0.641 [0.639;0.644] |

**Table S10.** Effects by chromosome and by telomere bin for each population.

**A**
| Chr | Rare |  |  |  | Non-synonymous |  |  |  | Neutral |  |  |  |
| --- | --- | --- | --- | --- | --- | --- | --- | --- | --- | --- | --- | --- |
|  | FCQ | EUR | ASN | AFR | FCQ | EUR | ASN | AFR | FCQ | EUR | ASN | AFR |
| 1 | **1.44** | **1.59** | **1.45** | **1.34** | **1.32** | 1.01 | 0.94 | 0.91 | **0.64** | **0.83** | 0.89 | 0.9 |
| 2 | 1.07 | **1.24** | **1.36** | 1.19 | 1.2 | 1.07 | 0.94 | 0.96 | **0.67** | **0.75** | 0.91 | 0.85 |
| 3 | **1.44** | **1.33** | **1.37** | **1.34** | **1.32** | 1.23 | 0.88 | 0.96 | **0.63** | **0.71** | 0.99 | 0.95 |
| 4 | 1.22 | 1.22 | **1.43** | **1.29** | 0.93 | 1.04 | 1.09 | 0.97 | 0.91 | 0.88 | 0.88 | 0.87 |
| 5 | **1.3** | **1.32** | 1.25 | **1.26** | **1.64** | **1.47** | **1.43** | 1.1 | **0.74** | **0.6** | **0.66** | **0.7** |
| 6 | **1.78** | **1.35** | 1.19 | **1.25** | **1.35** | 1.09 | 1.18 | 1.16 | **0.67** | 0.88 | **0.77** | 0.88 |
| 7 | 1.09 | **1.35** | **1.74** | **1.43** | 0.89 | 1.23 | 1.22 | 1.02 | 0.87 | **0.79** | **0.75** | 0.94 |
| 8 | **1.33** | **1.37** | **1.32** | **1.29** | **1.38** | 1.18 | 1.01 | 1.04 | **0.56** | **0.75** | 0.88 | 0.84 |
| 9 | **1.33** | **1.36** | **1.29** | **1.54** | **1.28** | **1.28** | **1.57** | **1.28** | **0.65** | **0.68** | **0.61** | **0.72** |
| 10 | 1.2 | 1.18 | 1.22 | **1.33** | **1.42** | **1.66** | **1.27** | **1.47** | **0.6** | **0.61** | **0.67** | **0.61** |
| 11 | **1.47** | **1.44** | **1.56** | **1.23** | 1.11 | **1.72** | **1.56** | **1.46** | **0.62** | **0.62** | **0.77** | **0.76** |
| 12 | **1.31** | **1.35** | **1.35** | **1.42** | 0.94 | 1.03 | 0.94 | 0.98 | **0.81** | 0.89 | 0.87 | 0.91 |
| 13 | 0.96 | **1.6** | 1.27 | 1.16 | 1.27 | 1.2 | 1.3 | 1.22 | 1.04 | 0.88 | 0.72 | 0.77 |
| 14 | **1.34** | **1.28** | **1.54** | 1.18 | 1.23 | 0.9 | 1.08 | 0.87 | **0.66** | 0.96 | 0.85 | 1.06 |
| 15 | 0.92 | **1.53** | 1.21 | 1.22 | **1.46** | 1.24 | 1.16 | **1.4** | **0.66** | **0.69** | **0.67** | **0.59** |
| 16 | 1.19 | **1.35** | **1.84** | **1.32** | **1.25** | 1.01 | 1.16 | 0.9 | **0.69** | 0.85 | **0.78** | 0.94 |
| 17 | **1.48** | **1.45** | **1.6** | **1.29** | 1.09 | 0.94 | 1.09 | 0.95 | **0.72** | **0.81** | **0.79** | **0.83** |
| 18 | 1.32 | 1.03 | 1.34 | 0.99 | **2.01** | 1.15 | 1.17 | 1.02 | **0.44** | 0.8 | 0.75 | 1.02 |
| 19 | **1.55** | **1.47** | **1.33** | **1.48** | 0.91 | 0.93 | 0.89 | **0.76** | 1.01 | 1.04 | 1.04 | **1.17** |
| 20 | **1.74** | **2.19** | **1.53** | **1.53** | **1.4** | 1.17 | 1.07 | 1.07 | **0.53** | **0.65** | **0.71** | 0.86 |
| 21 | 0.88 | **0.58** | 1.26 | 0.88 | **1.49** | 1.29 | 1.14 | **1.6** | **0.52** | 0.83 | 0.78 | **0.71** |
| 22 | 1.08 | 1.14 | 1.06 | **1.33** | 1.13 | 0.95 | **1.32** | 0.93 | 0.87 | 0.98 | **0.76** | 0.99 |

**B**
| Telomere bin<sup>a</sup> | Rare |  |  |  | Non-synonymous |  |  |  | Neutral |  |  |  |
| --- | --- | --- | --- | --- | --- | --- | --- | --- | --- | --- | --- | --- |
|  | FCQ | EUR | ASN | AFR | FCQ | EUR | ASN | AFR | FCQ | EUR | ASN | AFR |
| 1 | **1.35** | **1.29** | **1.16** | **1.2** | 0.98 | 0.84 | 0.99 | 0.69 | 0.83 | 0.96 | 0.85 | 1.31 |
| 2 | **1.24** | **1.37** | **1.41** | **1.26** | 1.06 | **1.54** | **1.47** | **1.27** | 0.94 | **0.73** | **0.79** | **0.86** |
| 3 | **1.36** | **1.45** | **1.27** | **1.28** | 0.92 | 1.12 | **1.15** | 1.09 | **0.85** | **0.81** | **0.81** | **0.84** |
| 4 | **1.45** | **1.6** | **1.51** | **1.45** | **1.39** | **1.21** | **1.2** | 1.03 | **0.56** | **0.74** | **0.73** | **0.88** |
| 5 | **2.17** | **1.47** | **1.42** | **1.48** | 1.1 | 1.04 | 0.99 | 0.86 | **0.77** | **0.78** | 0.87 | **0.93** |
| 6 | **1.29** | **1.24** | **1.29** | **1.18** | **1.49** | **1.24** | 1.11 | 1.09 | **0.62** | **0.78** | **0.78** | **0.87** |
| 7 | **1.12** | **1.28** | **1.5** | **1.16** | **1.28** | 1.08 | 1.07 | 1.02 | **0.62** | **0.78** | **0.79** | **0.83** |
| 8 | **1.18** | **1.26** | **1.38** | **1.4** | **1.16** | 1.12 | 1.03 | 1.05 | **0.74** | **0.76** | 0.86 | **0.8** |
| 9 | **1.31** | **1.35** | **1.33** | **1.34** | **1.21** | 1.02 | 0.94 | 0.94 | **0.72** | **0.88** | 0.92 | **0.88** |
| 10 | **1.15** | **1.22** | **1.44** | **1.2** | **1.33** | 1.09 | 1.19 | 1.08 | **0.72** | **0.83** | **0.79** | **0.88** |

<sup>a</sup>Each chromosome is divided in 10 bins of equal length, with bin 1 closer to centromere and 10 closer to telomere. Values in bold are significant OR. No significant differences were found, since permuting chromosome and telomere bin labels 1000 times lead to differences between chromosomes or telomere bins as extreme as seen in the data. However, the pattern on chromosome 19 is notable, as it shows a reverse effect compared to other chromosomes/genome-wide effects for non-synonymous and neutral variants in all populations.

